# How Obstacles Perturb Population Fronts and Alter their Genetic Structure

**DOI:** 10.1101/021964

**Authors:** Wolfram Möebius, Andrew W. Murray, David R. Nelson

## Abstract

As populations spread into new territory, environmental heterogeneities can shape the population front and genetic composition. We study here the effect of one important building block of inhomogeneous environments, compact obstacles. With a combination of experiments, theory, and simulation, we show how isolated obstacles both create long-lived distortions of the front shape and amplify the effect of genetic drift.

A system of bacteriophage T7 spreading on a spatially heterogeneous *Escherichia coli* lawn serves as an experimental model system to study population expansions. Using an inkjet printer, we create well-defined replicates of the lawn and quantitatively study the population expansion manifested in plaque growth. The transient perturbations of the plaque boundary found in the experiments are well described by a model in which the front moves with constant speed. Independent of the precise details of the expansion, we show that obstacles create a kink in the front that persists over large distances and is insensitive to the details of the obstacle’s shape. The small deviations between experimental findings and the predictions of the constant speed model can be understood with a more general reaction-diffusion model, which reduces to the constant speed model when the obstacle size is large compared to the front width. Using this framework, we demonstrate that frontier alleles that just graze the side of an isolated obstacle increase in abundance, a phenomenon we call ‘geometry-enhanced genetic drift’, complementary to the founder effect associated with spatial bottlenecks. Bacterial range expansions around nutrient-poor barriers and stochastic simulations confirm this prediction, the latter highlight as well the effect of the obstacle on the genealogy of individuals at the front.

We argue that related ideas and experimental techniques are applicable to a wide variety of more complex environments, leading to a better understanding of how environmental heterogeneities affect population range expansions.

**Author Summary:** Geographical structure influences the dynamics of the expansion of populations into new habitats and the relative importance of the evolutionary forces of mutation, selection, genetic drift, and gene flow. While populations often spread and evolve in highly complex environments, simplified scenarios allow one to uncover the important factors determining a population front’s shape and a population’s genetic composition. Here, we follow this approach using a combination of experiments, theory, and simulations.

Specifically, we use an inkjet printer to create well-defined bacterial patterns on which a population of bacteriophage expands and and characterize the transient perturbations in the population front caused by individual obstacles. A theoretical understanding allows us to make predictions for more general obstacles than those investigated experimentally. We use stochastic simulations and experimental expansions of bacterial populations to show that the population front dynamics is closely linked to changes in the genetic structure of population fronts. We anticipate that our findings will lead to understanding of how a wide class of environmental structures influences spreading populations and their genetic composition.

## Introduction

Populations expand into new territory on all length and time scales. Examples include the migration of humans out of Africa [1], the recent invasion of cane toads in Australia [2], and the growth of colonies of microbes. Although populations often persist long after invading [3], events during their spread can have long-lasting effects on their genetic diversity [4, 5]. Considerable effort has been undertaken to understand the role of the invasion process on the evolutionary path of the population: The small population size at the edge of the advancing population wave amplifies genetic drift, reducing genetic diversity, which can culminate in the formation of monoclonal regions [4]. The fate of mutations - deleterious, neutral, or beneficial - occurring in the course of the expansion depends on the location of their appearance with respect to the edge of the wave [6–8]. While the genetic consequences of such range expansions have been studied empirically [9, 10], the complexity of natural populations makes it difficult to draw general conclusions. Laboratory expansions of microbes have thus become a useful tool to illustrate, test and inspire theoretical predictions [11–14].

The majority of theoretical and experimental work on range expansions has focused on homogeneous environments while habitats in nature are often spatially heterogeneous with regard to dispersal or population growth, the two processes that lead to the expansion. Incorporating environmental heterogeneity into models of spreading populations [3, 4, 15, 16] raises complex problems. Heterogeneity can affect any parameter that controls population dispersal or growth and there can be many spatial patterns of heterogeneity. Ecologists and population geneticists often focus on different consequences of environmental heterogeneity. Work in population dynamics and ecology typically concentrates on the effect of heterogeneity on invasibility and the speed and impact of an invasion in such environments [3, 15, 17–20]. In contrast, population genetics studies usually assume a successful invasion and ask how environmental heterogeneities affect the population’s genetic composition [4]. Although heterogeneous carrying capacities [21], fragmented environments [22], single corridors or obstacles [23], and environmental patterns found on earth [24] have been addressed from a theoretical perspective, a systematic understanding is still missing. In this work, we study the population dynamics and relate the dynamics of the population front to the consequences on the genetic composition of the spreading population, thereby linking the evolutionary and ecological consequences of range expansions around obstacles.

We study the effect of isolated obstacles on the spread of populations, a scenario which is both complementary to a more commonly studied regime of isolated islands and forms the basis for studies of more complex environments consisting of multiple obstacles. What constitutes an obstacle or a barrier to the expansion depends on the population: For a macroscopic expansion of a terrestrial animal or a plant, lakes and mountain ranges are examples of obstacles. For microbes, regions with poor nutrients may represent an obstacle to colony growth. Using a combination of experiment, theory, and simulation, we characterize the obstacle’s effect in a regime of sizes where the shape of the front is well-described by a phenomenological model of expansion with constant speed. The constant speed model reveals general effects which hold independently of the mechanisms for population spread: The perturbation in the population front induced by the obstacle is determined by the obstacle’s width, but not by its precise shape. The front shape, induced by the obstacle, governs the effect on the genetic composition of the expanding population. Expanding past obstacles reduces genetic diversity and privileges alleles that just miss an obstacle’s edges, an example of ‘geometry-enhanced genetic drift’. This goes along with a characteristic change in the ancestry of individuals at the front which can be understood using the front’s dynamics. In addition to the phenomenological model of front shape, we study a reaction-diffusion model which enables us to compare experiments and model in more detail and to understand the utility of the constant speed in situations that extend beyond the experimental system studied here.

To derive these findings, we combine an analytical model, simulations, and experiments. While the experiments are the basis for theoretical work, they also allow us to test theoretical predictions. The analytical model provides the opportunity to make predictions for a variety of environments and length scales while simulations are used to explore regimes not accessible to analytical solutions. In addition to using established theoretical and experimental methods to study expanding populations, we present a new laboratory model system which allows us to quantitatively study population spread in heterogeneous environments: the expansion of bacterial viruses (bacteriophage) on a lawn of sensitive and resistant bacteria. Patches of resistant bacteria represent obstacles to the spread of the phage and can be generated using a printing technique, allowing us to quantitatively test predictions and offering the prospect of studying demographic and evolutionary processes in more complex, yet well-defined environments.

## Results

### Heterogeneous environments with binary growth conditions

We want to understand what happens when expanding populations confront environmental heterogeneities. For simplicity, assume that at each point the environment is a high quality habitat (large growth rate at population density well below carrying capacity) or a low quality habitat (very small or zero growth rate). If *ρ* is the fraction of the environment that allows growth, we can distinguish between two scenarios: For 0 *< ρ ≪* 1*/*2, the ‘island scenario’, a largely inhospitable environment is interrupted by islands or oases of growth; in contrast, for 1*/*2 ≪ *ρ <* 1, the ‘lake scenario’, a largely hospitable environment is punctuated by obstacles that impede growth (Fig. 1A). The island scenario, reminiscent of stepping stone models of population genetics [25], with a weak coupling between nearby islands by migration, is a situation where genetic drift can lead to genetic uniformity on individual islands due to founder effects [26] and has been extensively studied. Here, we address the generally less well studied lake scenario in the context of spatial expansions.

**Figure 1.**
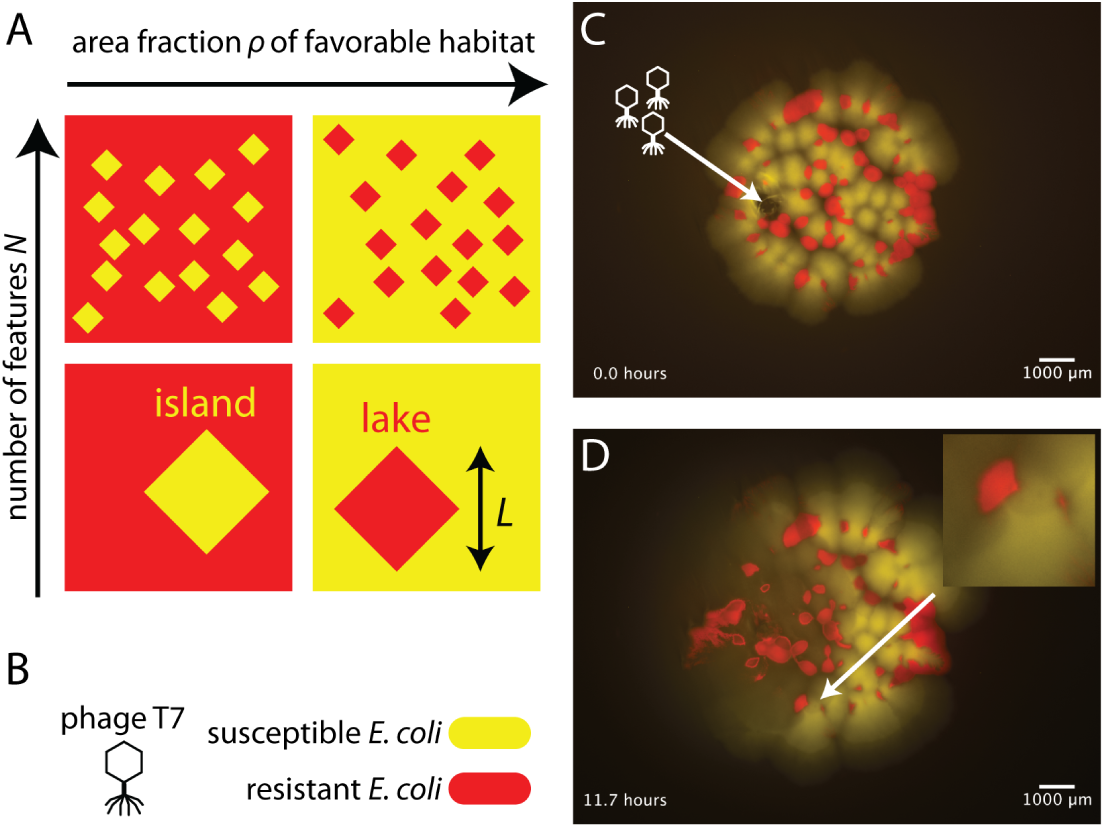
Plaque growth on a heterogeneous bacterial lawn. **(A)** Classification of environments composed of regions that permit or prohibit reproduction based on the area fraction of favorable habitat *ρ* and number *N* of features of linear size *L*. The features are embedded in the environment accessible to the spreading population. In this work, we focus on the ‘lake scenario’, i.e., regions that prohibit growth (red) distributed in an environment that permits growth (yellow). **(B)** Bacteriophage as an experimental model for expansion in heterogeneous environments: for bacteriophage T7, a lawn of susceptible *E. coli* (wild-type, WT) represents an environment of good growth conditions (yellow fluorescent marker), while a region with resistant *E. coli* (*waaC*Δ, red fluorescent marker) represents poor growth conditions. **(C)** Bacteriophage innoculation on a heterogeneous lawn. After inoculation with a carrier fluid that evaporates quickly, a mixture of susceptible and resistant cells grows into micro-colonies representing an environment with a large number of obstacles to phage growth (similar to A, upper right). Right before the colony was imaged, phage T7 was inoculated on the left part of the lawn, at the location indicated by the arrow. (The black gaps inside the lawn represent the absence of bacteria and are filled in in the course of the experiment.) **(D)** Bacteriophage spread on a heterogeneous lawn. After approximately 12 hours, the plaque (dark region due to lysis of bacteria) has extended through about half of the colony, almost exclusively affecting the susceptible part of the lawn (yellow). The regions of resistant bacteria cause transient perturbations in the front of the phage population, as seen in the close-up of the region indicated with a gray arrow. See S1 Video and S2 Video for more information.

In addition to the fraction of the habitat not occupied by obstacles, *ρ*, we must also consider the number of obstacles, *N*, in the new habitat to be invaded. When *N ≫* 1, i.e., when many (non-overlapping) obstacles are engulfed as the range expansion progresses, we expect that the principal effect of interest from a population dynamics perspective is that overall expansion slower than it would be without the obstacles (top of Fig. 1A). If, at the other extreme, the expansion domain only includes one obstacle, its size, shape, and spatial arrangement is expected to greatly influence the shape of the front on the length scale of the size of the habitat invaded (bottom of Fig. 1A). Below, we demonstrate that in addition the linear feature size of obstacles, *L*, governs their effect on the population front at the scale of individual obstacles.

### Transient perturbations of plaque boundary by obstacles

To explore the effects of obstacles on the population front dynamics, we employed a microbial model system, bacteriophage T7 spreading on a lawn of *E. coli* cells. Phage T7, a virus of *E. coli* infects bacterial cells and lyses them, releasing a large number of new phage particles which undergo passive dispersal and can infect nearby cells, a cycle of growth and replication that leads to an advancing population front. Phage T7 must kill the bacteria it infects [27] and its spread on a bacterial lawn is revealed by the growth of plaques (clearings in the lawn due to lysis of bacteria), a fast process easily visualized by bright-field or fluorescence microscopy. We produce a heterogeneous environment for phage spread by incorporating regions which do not support propagation of the population wave: While wild-type bacterial regions (marked by a constitutively expressed yellow fluorescent protein) correspond to regions supporting phage production, resistant bacterial patches (similarly marked by a red fluorescent protein (mCherry)) do not, see Fig. 1B and Materials & Methods. This lawn, with regions of susceptible and resistant bacteria, represents a static, heterogeneous environment that the phage population travels across during its expansion and that can be easily visualized.

Initially, we produced randomly distributed heterogeneity by inoculating a droplet of a dilute mixture of susceptible and resistant *E. coli* onto an agar plate. After inoculation, each cell grows into a microcolony, resulting in a bacterial lawn a few mm in diameter that consists of patches of susceptible or resistant bacteria (Fig. 1C). When we inoculated a drop of phage T7 at a discrete location, we saw the phage population spread on this heterogeneous lawn, with repeated cycles of infection and lysis of the susceptible bacteria leading to the loss of fluorescence and the expanding dark regions as shown in Fig. 1D. Approximately 12 hours after inoculating the phage, the plaque has extended halfway through the bacterial lawn (S1 Video). The phage front leaves behind a less-fluorescent region of debris, through which the patches of resistant bacteria expand further (compare Figs. 1C and D). The front shows several undulations, which are due to patches of resistant bacteria (“obstacles”) since these perturbations are absent in the control experiment lacking the resistant cells (S2 Video). Specifically, kinks in the front form behind obstacles as depicted at higher magnification in Fig. 1D. The overall speed of the plaque front is determined by additional parameters, which likely include the density of bacterial cells and their metabolic state: the front moves faster at the rim of the lawn both in the presence and absence of obstacles (S1 Video and S2 Video).

Fig. 2A displays the dynamics of the plaque boundary from Fig. 1D in more detail. A mostly linear population wave of phage encounters the region of resistant bacteria, an obstacle. The front curves as it passes the widest part of the obstacle and the two curved regions move along the far side of the object until they unite with each other, giving rise to a kink that disappears with time as the front moves beyond the obstacle. While the perturbation of the front by the obstacle and the formation of a lagging region is intuitive at first, we aimed for a quantitative model which can describe front shape and make predictions which can be tested experimentally.

**Figure 2.**
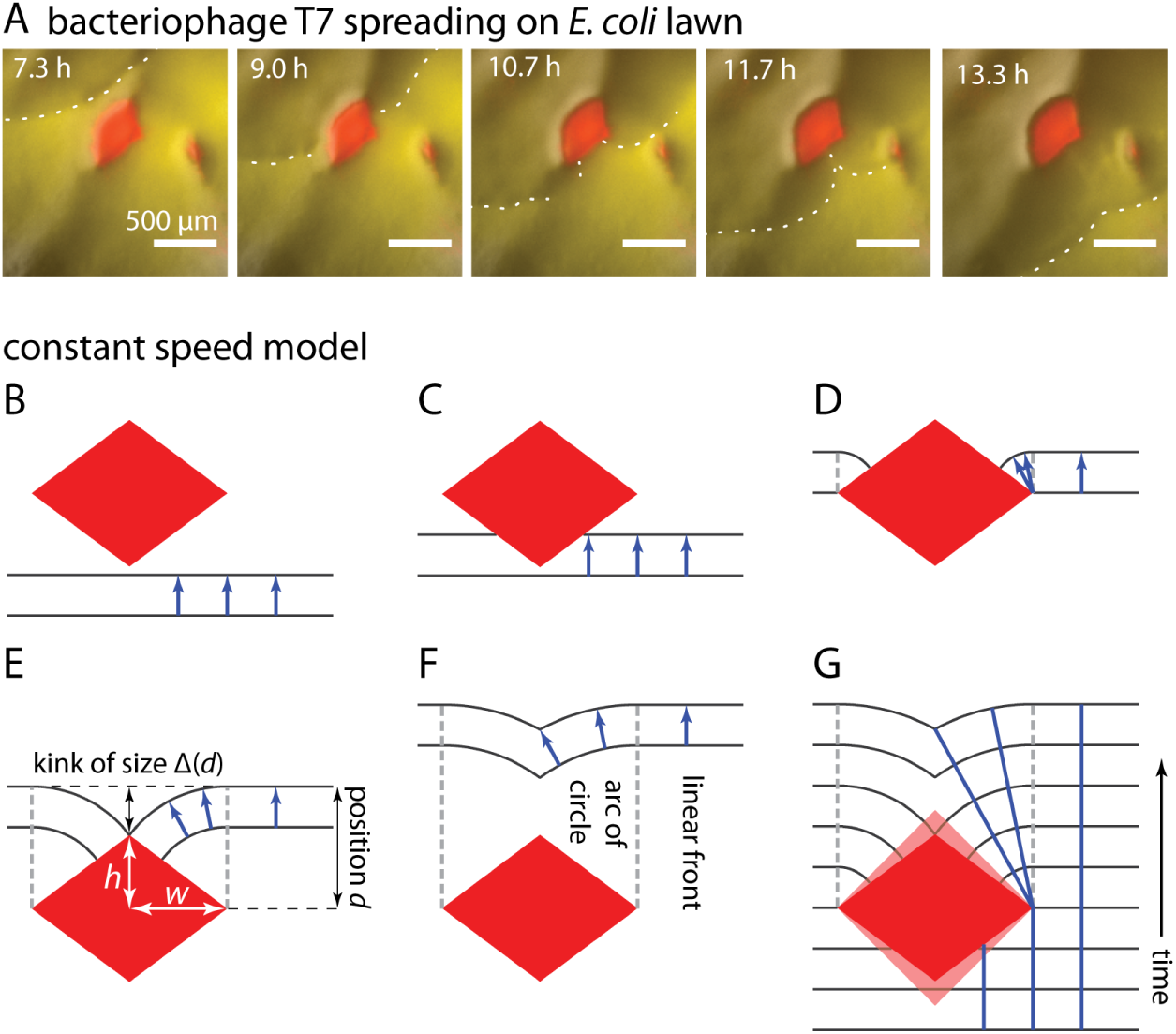
Formation of kinks behind obstacles and constant speed model. **(A)** Snapshots of plaque expansion (dark area) on lawn of susceptible bacteria (yellow area) around patch of resistant bacteria (red area). The region displayed is the region highlighted in Fig. 1D. The fluorescent images have been superimposed on a brightfield image to enhance visibility of the front, white dots have been manually included as guide to the eye. **(B-F)** Constructing the front shape encountering a rhombus-shaped obstacle using the constant speed model. Blue arrows have equal length and indicate the distance and direction of front movement over small time interval. Panels B to D illustrate how the front is broken as it encounters the obstacle and then curves around on the far side of the obstacle. Panels E to F show how a kink of size Δ, which forms when the disrupted front reunites, heals downstream from the obstacle due to the radius of the circular region that defines the perturbed front increasing with position *d* as the front moves beyond the obstacle. **(G)** Superimposition of the fronts in panels B to F (compare black lines in Fig. 4A,C) with blue lines indicating the path of a virtual marker at the front. For a lightly shaded rhombus with same width, larger height, the shape of the front is unchanged (but the circular arcs are shorter and the kink forms later).

### Constant speed model

An arguably most minimal model assumes that the front moves with constant speed in direction normal to the front and ignores the microscopic details of how phage replicate inside bacteria and diffuse outside them. We dubbed this the ‘constant speed model’. Fig. 2B-F illustrates the dynamics of a front encountering a rhombus-shaped obstacle: (B) An initially linear front moves forward uniformly until the obstacle is encountered. (C) When it encounters the obstacle, the front stays linear, but is interrupted in the interval where it would overlap with the obstacle. (This is different from scenarios where a front of material encounters an obstacle and the obstacle “pushes” the material to the sides.) (D) Beyond the obstacle’s widest points, propagation with constant speed creates circular arcs in the shade of the obstacle that are connected to a linear front on either side of the obstacle. The circular elements span a region given by the obstacle’s width and encounter the obstacle at a 90°angle. (E) For a rhombus with height 2*h* and width 2*w*, the arcs from the two sides meet and a kink forms after the front has traveled a distance 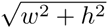 beyond the point of maximum width. (F) The kink then heals due to the increasing radii of the circular segments, i.e., the size of the indent Δ decreases (Δ(*d*) *∼* 1*/d* for large “downstream” distances *d*, where *d* is the distance, perpendicular to the front from the widest point of the obstacle to the unperturbed portion of the front, see below). Fig. 2G shows that the height of the rhombic obstacle does not play a role in determining front shape and thus the size of the indent while the kink heals: For an obstacle which is taller (light red rhombus), the kink forms later and the circular arc where it forms is correspondingly shorter, but the shape of the downstream kink is independent of the obstacle height. Moreover, the circular shapes of the front on the far side of rhombus-shaped obstacles all fall onto the same master curve when plotted in units of *w*. A calculation shows that the indent size Δ as function of position *d* is indeed independent of *h* and shows the same functional behavior if all lengths are expressed in units of *w* (see also the S1 Appendix):

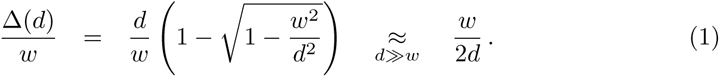

Counterintuitively, the width of the obstacle thus is a more important predictor of downstream front shape than obstacle height. For rhombus-shaped obstacles, obstacle height determines where the kink forms, but not the shape of the front after formation of the kink. Both *w* and *h* determine the linear feature size *L* introduced above (Fig. 1A). Below, we will discuss more general obstacle shapes and the influence of *L* on the applicability of the constant speed model.

#### Testing the constant speed model using printed environments

To test the predictions of the constant speed model, we designed an assay that allowed linear fronts of expanding phage populations to encounter obstacles of defined shape. We modified a method that used a consumer inkjet printer to print sugar solutions [28] to deposit bacteria in defined patterns on agar surfaces (such as was done using custom-made equipment [29]). The printer produces a field of bacteria on a rectangular (3.5 *×* 2cm^2^) agar patch at sub-mm resolution (Fig. 3A, Materials & Methods, S1 Protocol & S1 Fig). The printed founder cells grow into a lawn, which is inoculated with a linear front of phage T7 close to or at the region with resistant bacteria (Fig. 3A) and the front’s propagation is detected by fluorescence microscopy (Materials & Methods). Fig. 3B shows such a printed pattern at different stages of the phage invasion (see also S3 Video).

**Figure 3.**
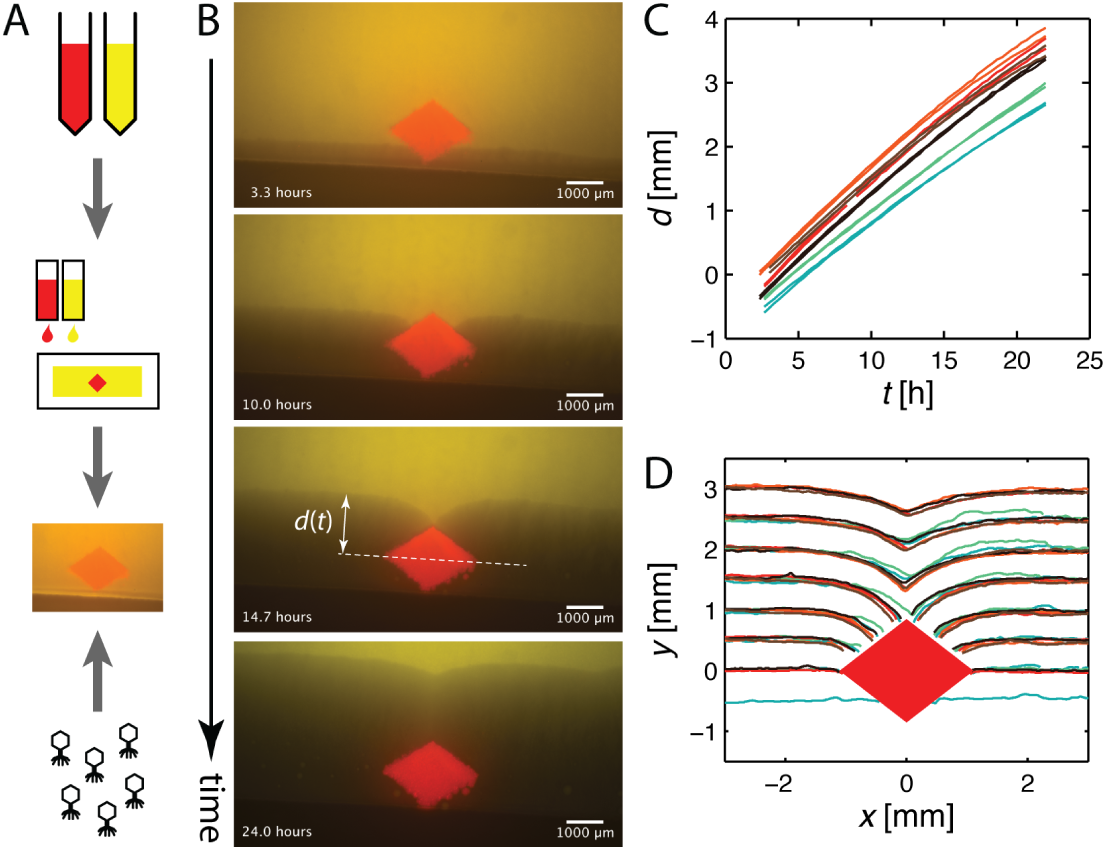
Printing bacterial lawns and plaque growth in well-defined environments. **(A)** Schematic diagram of the assay to observe plaque growth in well-defined reproducible environments. A digital representation serves as input for printing bacterials strains, both wild-type and phage-resistant, on an agar patch using a consumer inkjet printer. After the pattern has grown, phage is added using a piece of membrane impregnated with a phage solution and plaque growth is observed. See Materials & Methods for details. **(B)** Snapshots of plaque propagation (dark regions) around a rhombus-shaped area of resistant bacteria (red) printed in a sea of sensitive bacteria (yellow). The plaque front remains flat until it reaches the widest part of the obstacle. There, it curves into a region roughly as wide as the obstacle. Once the front reaches the top of the obstacle, a kink forms, which then slowly heals. Panel (B) also illustrates *d*(*t*), the distance the front travels beyond the obstacle at time *t*, where *d* = 0 at the point of maximal width of the obstacle. **(C)** Position of the front relative to the obstacle center as inferred by quantitative image analysis (Materials & Methods), as a function of time, for six technical and biological replicates. Two lines of the same color indicate the front position determined 3 mm to left and right of the obstacle center, respectively. The velocity is roughly constant, although a gradual slow-down is noticeable. **(D)** Front shape at different front positions *d* relative to the obstacle for six technical and biological replicates together with shape of the obstacle as inferred by image analysis (Materials & Methods), colors corresponding to panel (C).

We used the difference between two consecutive images to detect the front of the plaque (Materials & Methods), and studied the front position as a function of time. Similar to the above discussion, we define the unperturbed front position *d*(*t*) as position of the plaque’s edge at a horizontal distance of *±*3 mm away from the obstacle center. Fig. 3C shows that the plaque grows at an approximately constant speed, but slows down slightly over time, presumably due to *E. coli* entering stationary phase [30]. The varying slope illustrates variation in front speed among replicates. Overall, the plaque extends with an approximately constant speed of 0.2mm*/*h. The profile of the fluorescence signal perpendicular to the front is constant as shown in S2 Fig. Fig. 3D shows the front shape over time for multiple replicates. The evolution of the front is very similar across replicates, despite small variations in front speed and initial conditions, and agrees qualitatively with the constant speed prediction of Fig. 2G.

The constant speed model predicts the shape of the front, scaled by its width, to be identical for all rhombus-shaped obstacles. To test this prediction, we repeated the experiment for four different obstacles, combining two different widths with two different heights. Figs. 4A,B display front shapes and the indent size as measured for all four (colored) obstacle shapes. As predicted, the data collapse very well onto single lines if lengths are divided by *w*. (This is not the case for other scalings, see S3 Fig). Although the constant speed model successfully predicts how the front shape scales with the obstacle’s width and the front’s distance from the obstacle, the details of the predicted front (black line) differ from the experimental data. The experimental profiles consistently lag behind the predicted front and, equivalently, have a larger indent size than predicted (Figs. 4A,B). However, this quantitative disagreement does not affect the scaling behavior of Δ(*d*) (Fig. 4B, see S4 Fig for the same data on linear scale).

**Figure 4.**
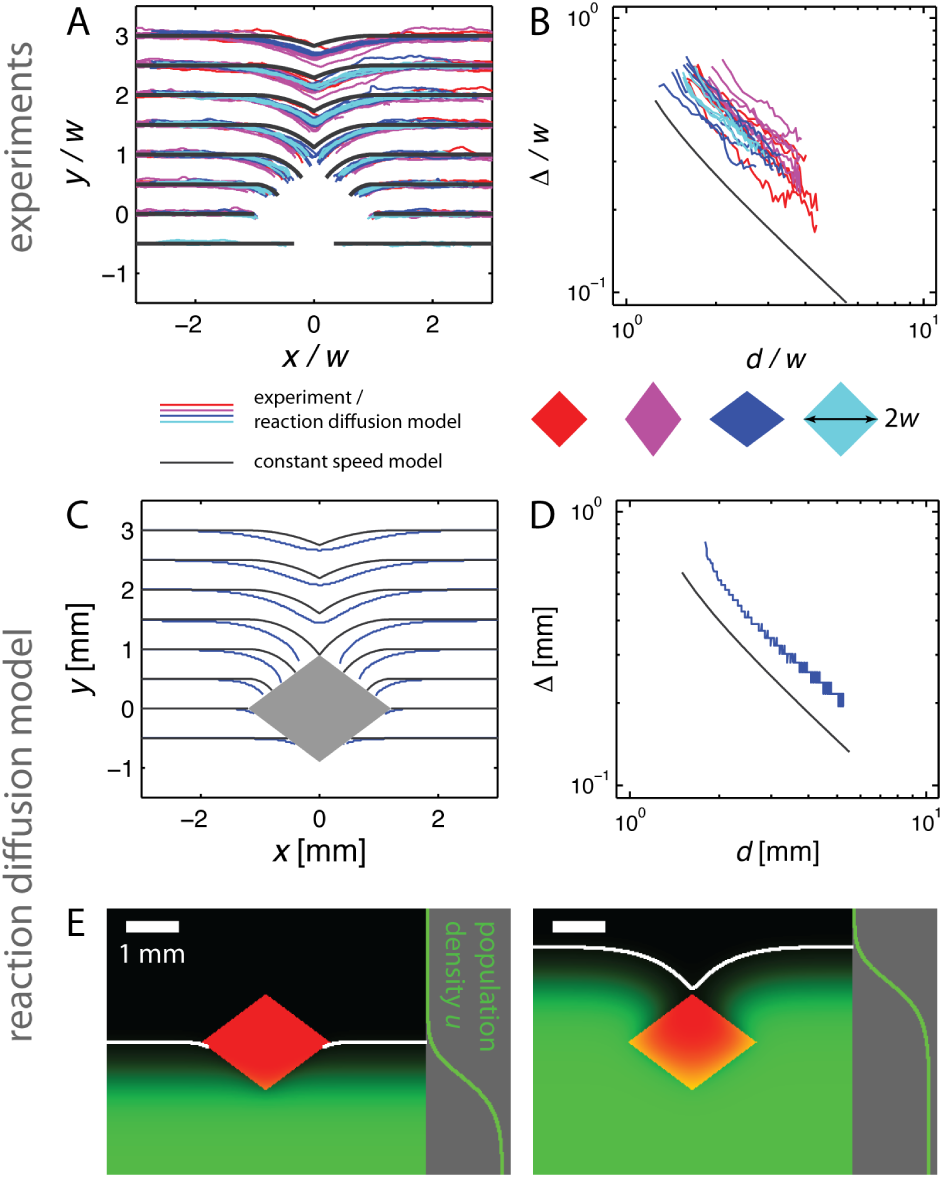
Comparison of phage fronts and reaction diffusion model to constant speed model predictions. **(A)** Experimentally determined plaque front shapes (*x, y*) for different obstacles with length in units of *w*, half the obstacle width (colored shapes, colored lines). While the data collapse as predicted by the constant speed model, a lag of the experimentally measured fronts is apparent relative to the front shape predicted by constant speed model (black line). **(B)** Data from the same experiments as shown in (A), but plotting the indent size Δ as function of front position *d* on a logarithmic scale (see Fig. 2E). The slope of the data is consistent with the black line, which is the prediction (Eq. 1) of the constant speed model. **(C-D)** Comparison of the front shape (C) and indent size (D) between the numerical solution of the reaction-diffusion model (colored lines) and prediction of the constant speed model (black lines) for one obstacle. A lag, such as the one observed experimentally, is clearly visible, see also S5 Fig. **(E)** Snapshot of the numerical solution of the reaction diffusion model. A traveling population wave encounters a region with zero growth (red rhombus). Population density is indicated in green, its profile far away from the obstacle is shown on the right. The white line indicates the inferred front. See Materials & Methods for details.

### Reaction-diffusion modeling rationalizes limitations of constant speed model

The constant speed model captures the general features of the front dynamics observed in the phage experiment, but the deviation prompted us to study a more detailed model which considers more of the details of phage propagation. In addition to understanding the deviation, the more detailed model will shed light on the range of applicability of the constant speed model allowing us to generalize beyond the phage system as we will show below.

The dynamics of plaque growth on *homogeneous* lawns has attracted considerable interest in the past [31–35]. A reaction-diffusion model, which captures the phage-bacterial interaction, the phage life cycle, and focuses on bacteriophage T7, has been suggested by Yin and McCaskill [32]: phage bind bacteria to form infected cells, and these, with a rate constant, burst to release more phage. More complex successor models focusing on the delay between infection and release of progeny phage have been published [34]. We decided not to generalize these models to heterogeneous environments for two reasons: (i) The appropriate parameters are not known for our experiments and (ii) we want a general description that allows us to reach conclusions that extend beyond the infection of *E. coli* by bacteriophage T7.

We therefore employed a coarse-grained reaction-diffusion model in which a species disperses by diffusion and replicates locally with logistic growth (the local reproduction rate increases linearly with population density, then decreases and reaches zero at the carrying capacity of the environment) except inside of obstacles. In the absence of obstacles, this model produces a traveling population wave with an exponentially shaped leading edge that moves at constant speed like the population wave resulting from the model by Yin and McCaskill [32]. Mathematically, it is a generalized version of the Fisher-Kolmogoroff-Petrovsky-Piscounoff equation (FKPP equation) [36–39], which captures the two processes underlying a range expansion, dispersal and growth. In our generalized model, the time evolution of viral population density *u*(**x***,t*) depending on location **x** and time *t* is given by:

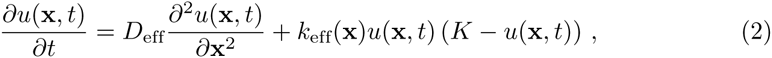

where the first term describes dispersal by diffusion with an effective diffusion coefficient *D*_eff_. The second term captures local logistic growth with reproductive rate *k*_eff_ and constant carrying capacity *K*. By rescaling the phage density *u*(**x***,t*), we can set *K* = 1 without loss of generality. In general, *k*_eff_ will depend on the bacterial density, the number of phage an infected bacterium releases (the burst size), the adsorption kinetics of the phage, the rate for lysis of infected host, etc. [32]. Inside the obstacle, we will set *k*_eff_ = 0.

We used our data to estimate the values for the phage’s diffusion coefficient, *D*_eff_, and reproductive rate, *k*_eff_. For biologically relevant initial conditions, an unimpeded, linear population front moves forward with front speed 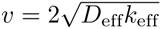 and front width parameter 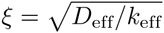 [39]. From Fig. 3C we find that the plaque front extends with a speed of about 0.2mm*/*h. With a rough estimate of the diffusion coefficient of 0.0144 mm^2^*/*h ([32, 40], see Materials & Methods) we can determine an effective growth rate of *k*_0_ = 0.7 */*h for the phage in our experiments. We assume that the phage’s diffusion coefficient inside the obstacle remains the same, but that no growth is possible due to the lack of susceptible bacteria, thus allowing us to set *k*_eff_(**x**) = 0 inside the obstacle and *k*_eff_(**x**) = *k*_0_ otherwise.

We next numerically solved Eq. 2 for the four different obstacles considered experimentally. Figs. 4C,D display the fronts and indent size, while Fig. 4E displays two snapshots from the numerical solution of the wide obstacle (see S4 Video). To quantify front shape at the leading edge, we defined front position as the boundary at which the population density is larger than 5 % of the carrying capacity (white line in Fig. 4E, see Materials & Methods). For the wide obstacle (and the three other obstacles, S5 Fig) we observe a lag of the front for the numerical solution (colored line) relative to the constant speed prediction (black line), in qualitative agreement with the experiments. This lag also manifests itself in an increased indent size (Fig. 4D). To test sensitivity to the value of the diffusion coefficient, *D*_eff_, we repeated the analysis for the wide obstacle with *D*_eff_ *→* 3*D*_eff_ and *D*_eff_ *→ D*_eff_*/*3 and found the lag to persist in both cases (S6 Fig, Materials & Methods). As expected, for decreasing *D*_eff_ the lag, relative to the constant speed prediction, becomes smaller. Taken together, the numerical solution of the reaction-diffusion model produces a lag similar to that seen in experiments of the phage model system (Fig. 4) even though its parameters were not derived from the front’s shape.

Where does the lag originate from and under which circumstances is the constant speed model a good description? At short times and small length scales, diffusion dominates the evolution of the population density. Kinks in the front will eventually be rounded and small bulges in the front smoothed out. (We disregard possible number fluctuations at the frontier and associated possible front instabilities [41]). But at long times and large length scales, diffusion is slow compared to the deterministic process of invasion and thus the effects of diffusion are limited to small scales. This critical length dividing these two regimes is given by *D/v*, the ratio of the diffusion coefficient *D* to the speed of the advancing front *v*. Up to a prefactor, this ratio is given by the front width parameter 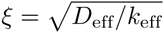 and is proportional to the width of the profile, perpendicular to the front, which is shown in Fig. 4E and S4 Video. We therefore expect the deterministic process of invasion to play the major role in determining front shape on length scales much larger than *ξ*, justifying the use of the constant speed model as an approximate, but intuitively useful model for understanding how population spread around obstacles. For our experimental system, *ξ ≈* 0.1 mm which is considerably, but not strikingly, smaller than the scale determining the shape of the obstacle (1 − 2 mm).

The simplicity of the reaction diffusion model (Eq. 2 only has two free parameters.) allows us to identify two mechanisms for the lag of the front relative to the constant speed model. First, virions diffuse into the obstacle, recognizable by the obstacle in Fig. 4E slowly turning yellow at its boundaries. This viral sink leads to a reduced population density close to the boundary and thus to a lagging boundary. This lagging region is limited to a boundary layer whose width is of the order of the front width parameter *ξ* (S7 Fig, panel A). The boundary layer moves at the same velocity as the unperturbed front, but can be an important constituent of the perturbed front. Because our obstacles are only one order of magnitude larger than *ξ*, we expect the boundary layer to contribute to overall front shape and thus to the observed lag. (We attribute the differing shapes of the front at the boundary layer between experiment (Fig. 4A) and theory (Fig. 4C) to the coarse-graining embodied in our model and differences in front detection). Second, the constant speed model predicts a very large curvature of the front close to the position of maximum width, which leads to a slowing of the front for radii of curvature smaller than *ξ* [39]. We therefore expect a contribution to a lagging front whenever the front of the obstacle is kinked or curved (S7 Fig, panel B). Both corrections should be small in the limit *L ≫ ξ* (S1 Appendix), meaning that the constant speed model can be used at large scales.

### On large scales, the coarse-grained model predicts universal front shape

Since we expect the constant speed model to successfully predict the front shape for large obstacles, we can construct the front shapes for more general obstacle shapes and infer general properties of front shape that are independent of the shape of the obstacle (see below and the S1 Appendix), which is not possible using experiments or numerical solutions alone.

While for rhombus-shaped obstacles the front shape is particularly simple (the front consists of two linear and two circular segments only, Fig. 2G), we now consider general convex, mirror-symmetric obstacles. When the front encounters an obstacle (as when it first envelops the tip of a rhombus or the front half of a circle), the shape of the front remains planar. As the obstacle starts to decrease in width, each point along the boundary is the source of a circular segment contributing to the front (similar to Fig. 2D) and the front thus encounters the obstacle at a 90°angle. Eventually, a kink or a “cusp” (a kink with infinite slope) forms on the far side (S8 Fig), which heals downstream from the obstacle. Note that this implies that changing the size of the obstacle (without changing its shape) implies that the front’s overall shape stays unchanged, but gets scaled by the same factor that the obstacle size increased or decreased.

In addition, as the kink heals downstream from the obstacle, we eventually recover a scaling result similar to Eq. 1. In this respect, the front exhibits a universal behavior far away from the obstacle: the perturbation inherited by the front is determined by the obstacle’s width, but not by its precise shape. Some quantitative predictions of the constant speed model for isolated circular, elliptical and elongated tilted obstacles are found in the S1 Appendix (S8 Fig, S9 Fig). For objects that are not convex, we expect a similar overall behavior. An obstacle with a complicated shape still results in a kink which gradually heals as the front moves beyond the obstacle (S10 Fig, S5 Video).

### Fate of alleles is determined by their location relative to the obstacle

We next examine how genetic composition of a population is shaped by obstacles it encounters, first predicting the obstacle’s effects based on the constant speed model followed by examining an experimental model system and simulations. As populations expand, genetic drift leads to the local reduction of genetic diversity. Thus we consider a population front that contains different neutral alleles at different positions along the front encountering an obstacle whose width is greater than the width of the front that each allele occupies.

Fig. 5A displays a series of front shapes together with a simplified set of initial allele distribution indicated by orange, green, cyan, blue, and red colors. In the spirit of the constant speed model, we focus on front shape dynamics alone and neglect sector boundary wandering. The front segment with the cyan allele either cannot propagate within the obstacle or, in the case of a virus like T7, slows down dramatically since only diffusive motion is possible. Hence, this allele is lost and does not contribute to the range expansion at later times. After passing the point of maximum width, the circular arcs of the front in the ‘shadow’ of the obstacle grow due to inflation and therefore alleles marked in green and blue occupy a larger fraction of the front. As the kink heals, the green and blue alleles occupy the part of the frontier that lies in the shadow of the obstacle. The abundance of these alleles, which were a small fraction of the initial front, stays elevated even after the kink has healed. Note, however, that part of the increase in allele abundance is transient since the arc length of the circular segments gets reduced during healing of the kink, although the radius still grows and the front thus locally experiences inflation. Fig. 5A depicts a special symmetric initial condition of allele frequencies that guarantees that alleles benefitting from the inflation in the wake of the obstacle (green and blue alleles) will meet precisely at the top of the obstacle. However, selectively neutral, grazing alleles will meet at the top for quite general initial conditions, i.e., there is always a sector boundary that gets ‘pinned’ at the top of the obstacle.

**Figure 5.**
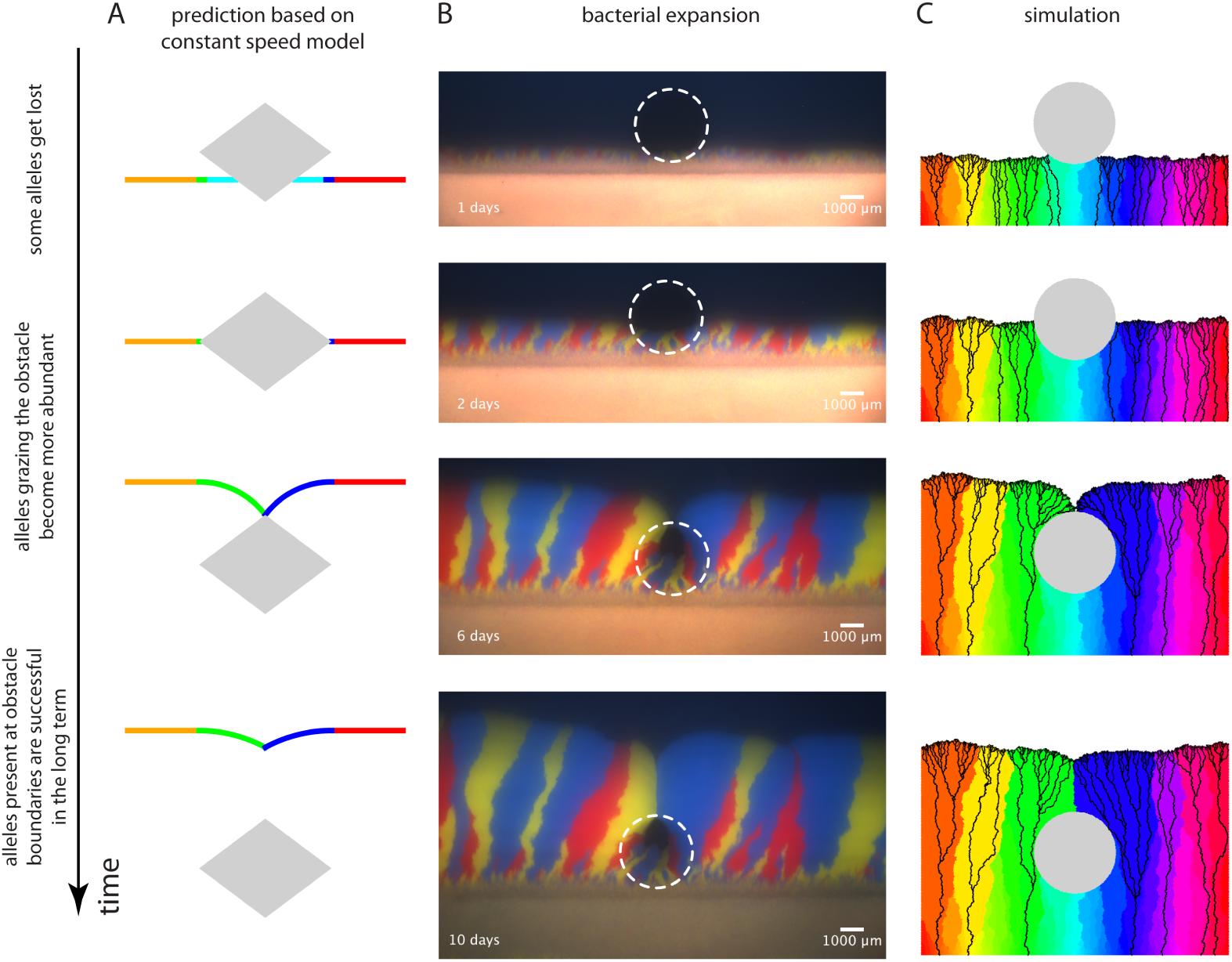
Effect of obstacle on allele abundance at front. **(A)** Illustration of the changes in allele abundance predicted by the constant velocity model. Alleles exactly at the frontier can be (i) driven to extinction (cyan), (ii) largely unaffected by the obstacle (orange & red lines), or (iii) sweep through a substantial part of the population after grazing the corners of the obstacle (green & blue lines). The linear region of increased abundance decreases somewhat as the kink heals. **(B)** Spatial expansion of *E. coli* with three different fluorescent labels on agar plates around a circular region with reduced nutrients (white dashed circle serves as guide to eye) after approximately 1, 2, 6, and 13 days of growth. In addition to the features outlined in the text, the sector boundary, where the two grazing sectors meet to form the kink in the population front, appears to be significantly more straight than the other sector boundaries in the colony. We speculate that this behavior emerges from a combination of geometrical effects and an increased local population size at the kink. Finally, for the replicate shown here, the sector boundary behind the obstacle is aligned in the propagation direction of the front but the majority of the remaining sector boundaries display a slight bias to the right. The tilt to the right is equivalent to the chirality of sector boundaries seen for some *E. coli* strains in circular spatial expansions [11, 42] and the straight boundary behind the obstacle suggests that something is phenomenologically different when two sectors meet each other. **(C)** Results of stochastic simulations of a lattice-based model with about 600 alleles encountering a circular obstacle (gray area). Black lines are lineages together representing the genealogy of individuals at the final front position. The individual experimental replicate and simulation were chosen to highlight the features described in the main text; other replicates and simulations show the same overall behavior and are consistent with the description in the main text.

The constant speed model argues that alleles that fail to encounter the obstacle will be unperturbed, those whose segment of the front entirely collides with the obstacle will be eliminated, and those that graze the obstacle will be privileged because they will fill in the region downstream of the obstacle.

We tested this idea experimentally by using fluorescent proteins as labels for selectively neutral alleles. Because we could not produce expansions with fluorescent phage, we used the expansion of three *E. coli* strains, which express different fluorescent proteins. Two of the strains have been characterized previously [11] and we constructed a third strain which behaves comparably for the purpose of the experiment. We created heterogeneous agar plates by adding a circular membrane with an impermeable region just below the top layer of agar. We then launched a linear expansion of a mixtures of the three marked strains and observed them as they grew past the circle that blocked access to nutrients (Materials & Methods, S11 Fig). Fig. 5B displays the range expansion after approximately 1, 6 and 13 days of growth (see S6 Video for additional time points and Materials & Methods for a description of replicate experiments).

Before the population meets the obstacle, genetic drift at the population front leads to separation into monoclonal regions of the three different colors [11, 43] allowing us to see the effect of the obstacle. The sectors that encounter the obstacle head on are lost but the two that just graze the obstacle grow in its shadow, increasing the abundance of the corresponding alleles, which meet at the top of the obstacle. These features are experimentally reproducible and verify the predictions of the constant speed model.

Next, we performed stochastic simulations, in which organisms reproduce on a lattice. Occupide sites on a lattice are copied onto unoccupied neighbors thus propagating the front [5] (a variant of the Eden model [44, 45] extended here to track genotypes, see S12 Fig and Materials & Methods). The obstacle is a set of lattice sites which cannot be occupied. Fig. 5C shows an initially linear front encountering an obstacle. The obstacle leads to dynamics that are qualitatively similar to the bacterial range expansion described above (S7 Video). Fig. 5C shows that individual alleles can go extinct by two processes: the wiggling of sector boundaries caused by genetic drift [5, 11, 43] and collision with the obstacle (light green to light blue sectors, Fig. 5C). The alleles that graze the corners that define the obstacle’s width dominate the curved part of the front during the subsequent inflationary phase (green and purple sectors, Fig. 5C) and meet at the top of the obstacle.

The founder effect of individuals near the point of maximum width also dominates the population’s genealogy downstream of the obstacle. Black lines in Fig. 5C represent lineages of individuals at the front. As already evident from the labeling of alleles with colors, none of the lineages pass through the area in front of the obstacle. In addition, most of the individuals at the curved part of the front originate from a small number of ancestors near the point of maximum width. Strikingly, none of these lineages pass through the point where the two populations meet behind the obstacle. Despite the expansion of the green and purple sectors right before they encounter each other, the parts of the population which meet at the top of the obstacle have no descendants at the front at late times. This effect arises because the two sectors encounter each other (almost) head on just behind the obstacle (Fig. 5C). Although this effect does not manifests itself in the sectoring pattern we deduced from the constant speed model (Fig. 5A), it can be understood within the framework of the model: In Fig. 2G, blue lines indicate the position of a virtual marker at the front coinciding with the overall shape of lineages behind the population front. This suggests that the constant speed model may also be used to predict the evolutionary dynamics of a spreading population in more complex environments.

In summary, we found that the constant speed model used to describe the front shape of a viral population expansion can be used to understand the effects of an obstacle on the diversity of neutral alleles in an expanding population. These include the loss of alleles encountering the obstacle head-on and a founder effect from individuals present at the point of maximum width. Since the obstacle does not affect fitness of individuals carrying specific alleles, but in an intricate way increases random fluctuations, these effects are an example of ‘geometry-enhanced genetic drift’.

## Discussion

Organisms rarely spread across featureless habitats. Instead, they must find ways to survive and reproduce in the presence of environments that are heterogeneous in space and time. To investigate the effects of spatial heterogeneities on the dynamics and genetics of a spreading population, we combined experimental and theoretical approaches to understand the effect of single obstacles, of defined geometry, where organisms could not reproduce. When bacteriophage T7 encounters resistant *E. coli* the bacteriophage population front is perturbed in the wake of the obstacles by a sharp kink that slowly heals as the front moves on. A constant speed model gives an intuitive understanding of this perturbation, and a more detailed, reaction-diffusion model rationalizes the deviation between experiment and the constant speed model’s predictions. In addition, the constant speed model explains that in a genetically diverse population, alleles that run into the obstacle are eliminated and those that graze its sides increase in abundance, an example of geometry-enhanced genetic drift.

A mathematically stringent analysis by Berestycki et al. predicted transient perturbations of planar waves encountering a single compact obstacle [46]. From a physical perspective, when the obstacle size *L* satisfies *L ≫ ξ*, where *ξ* characterizes the front width, considerable understanding of the perturbation is possible using a model based on front propagation locally and with constant speed. In this limit, the shape of the front can be found using a straightforward geometric construction that has an analogy in geometrical optics (S1 Appendix). Interestingly, in this regime, a linear front stays unperturbed while it envelops the obstacle, in contrast to a first intuition based on a front of fluid material encountering an obstacle. Our analysis of the front shows that the width of the obstacle, and not its precise shape, determines the long-term dynamics of the perturbation caused by the obstacle. Extensions to many obstacles should be possible in this regime. If obstacles are sparse, the healing of the perturbations implies that the front speed should be only marginally reduced compared to expansion in the absence of any obstacles. If obstacles are close enough to each other that the perturbation from the preceding obstacle has not healed much before the next obstacle is encountered, the perturbations will add up and an ensemble of obstacles will reduce front velocity measured perpendicular to the front. Obstacles regularly placed on a lattice are a special case: the existence of open channels, (unobstructed by obstacles and much wider than the front width parameter) will allow the front to travel as fast as it would without obstacles; the remaining territory will then be explored in the wake of the front. This approach will complement recent studies using reaction-diffusion models to study invasion in heterogeneous environments [18].

If the density of obstacles is so high that no free paths connect the different boundaries of the environment, the traveling wave cannot propagate around obstacles. When dispersal within obstacles is possible, the population can nevertheless expand via migration between regions with good growth conditions, which is essentially the island scenario depicted in Fig. 1A. Invasion is not possible in a scenario where population spread is hindered by a connected set of impermeable obstacles (compare to the percolation threshold concept [47]).

When the size of the obstacle approaches the parameter that sets the width of the population front, the constant speed model breaks down. This regime can be understood by numerically solving a two-dimensional reaction diffusion system (a generalized FKPP equation), which rationalizes the lag between the experimentally observed phage front and the constant speed model and bridges the gap to the regime where the length scale of the heterogeneities in the environment is much smaller than *ξ* and perturbations in front shape are therefore not expected.

When the obstacle strongly perturbs the shape of the population front, we predicted that these perturbations affect the fate of alleles and lead to ‘geometry-enhanced genetic drift’. Analyzing the fate of lineages shows that the descendents of individuals trapped in front of the obstacle or born right behind it are lost in the long term. Our results are in qualitative agreement with a simulation study that demonstrated a decreased probability of survival of neutral (and deleterious) mutations occurring just in front of and right behind an obstacle [23]. Taken together, our results show that the long-term reproductive success of an individual depends on its position relative to the obstacle the population encounters as well as the random sampling that drives genetic drift, expanding the list of factors that contribute to ‘survival of the luckiest’ [48]. Theory and simulation that include the occurrence and fixation of new mutations during range expansions around obstacles should illuminate the process of allopatric speciation during a range expansion.

Single obstacles could have pronounced effects on evolution beyond shaping the abundance of neutral alleles. Because the small subpopulation that grazes the obstacle expands spectacularly, obstacles could make it easier for deleterious mutations to survive. This expansion protects deleterious mutations from extinction [49] and could establish a subpopulation which is large enough to survive for a considerable amount of time. This time span might be long enough for a second, beneficial mutation to occur, which has implications for the crossing of fitness valleys. Because the obstacle is not a population bottleneck, failure to acquire such a second mutation does not lead to a reduced fitness of the population in the long-term: genotypes that passed further away from the obstacle would eventually spread sideways and extinguish the deleterious allele if it is not rescued by a second beneficial mutation. A true spatial bottleneck would have fixed the deleterious allele and thus reduced the (absolute) fitness of the whole population. The rapid evolution of phage should allow such questions to be addressed experimentally in the future.

The models we used to describe the spread of phage populations were successful, even though they ignored the details of the bacteriophage life cycle. We found that for large obstacles the constant speed model is a good description for the front shape and expect in consequence the effects of ‘geometry-enhanced genetic drift’ to hold. How do these results apply to organisms whose spreading mechanism is very complex or even not well characterized? Spread of a population is caused by the combined effects of dispersal and growth. Random dispersal, even active random motion, thereby is only relevant on short scales. We thus expect that for very large obstacles (relative to the scale at which dispersal ceases to be relevant) the constant speed model to be a good description. For a given species, one would therefore estimate effective diffusion and growth rates and compute the characteristic length 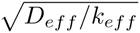. If this length is smaller than an obstacle, such as a lake or a mountain, the constant speed model is expected to be a useful null expectation for the spread of the population.

Similarly, although our analysis focused on single obstacles, we believe that our findings can be extended to natural environments, which typically display more complex heterogeneities. As a first step, by using the findings for isolated obstacles we expect to be able to describe observables such as the effective front speed and an ‘effective genetic drift’ in environments with obstacles such as those displayed in Fig. 1A.

## Materials & Methods

### Experimental Procedures

#### Plasmids

To fluorescently label bacteria, we used three different plasmids which (only) differed in the ORF for the fluorescent protein and the immediately surrounding nucleotides. The vector is pTrc99A, which provides resistance against ampicillin and expresses the *lac* repressor [50]. The ORF of the fluorescent protein is cloned between the SacI and XbaI sites and is under control of the *trc* promoter, a hybrid of the *trpB* and *lac* promoters. This promoter provides IPTG inducibility of the expression of the fluorescent protein. However, throughout our experiments, we did not add IPTG to the medium since we found the background expression to be sufficient for our experiments. Two of the plasmids (encoding for fluorescent proteins CFP and venus YFP) were identical to those described by Hallatschek et al. [11], while the third (encoding for mCherry) was obtained by substituting the ORF.

#### Strains *E. coli* - heterogeneous bacterial lawn

Two strains of *E. coli* were used to produce a heterogeneous lawn for bacteriophage T7. One strain, *E. coli* BW25113 (CGSC# 7636) is susceptible to T7 phage infection, the other strain is partially resistant by means of the deletion of the *waaC* gene (also known as the *rfaC* gene) [51] (JW3596-1, CGSC# 11805, part of the Keio collection [52]). (The product of the *waaC* gene is involved in the synthesis of the lipopolysaccharide whose recognition is essential for adsorption of the phage [51].) Both strains were obtained from the Coli Genetic Stock Center (CGSC, New Haven, CT). The susceptible strain was transformed with the plasmid expressing venus YFP (resulting in strain eWM43), the resistant strain was transformed with the plasmid expressing mCherry (resulting in strain eWM44).

#### *E. coli* - bacterial expansion

To study range expansions of bacteria with different alleles, three strains of *E. coli* DH5*α* transformed with the plasmids described above were used. Two of the strains were identical to those used by Hallatschek et al. [11] (expressing CFP (named strain eWM282) and expressing venus YFP (named strain eWM284)) and had been shown to be selectively neutral. The third strain was obtained by transforming *E. coli* DH5*α* with the plasmid expressing mCherry (eWM40) and showed a comparable front velocity when grown together with the two other strains (see Fig. 5B).

#### Bacteriophage T7

We originally obtained bacteriophage T7 as an aliquot from the wild-type stock of the Richardson laboratory (Harvard Medical School, Boston, MA). To obtain phage optimized for plaque spreading, phage was picked from the rim of a plaque growing on a top agar lawn of *E. coli* of the BW25113 background and subsequently grown in liquid *E. coli* culture of the BW25113 background. The lysate was mixed with NaCl to a final concentration of 1.4 M. The supernatant of the subsequent centrifugation step was stored at 4°C. Aliquots of this stock were used for all experiments.

#### Plaque growth in randomly structured heterogeneous environments

Plates (outer measure: 86 mm *×* 128 mm) were filled with 40 ml media (2xYT, 20 g*/*l agar and 100 *μ*g*/*ml ampicillin) and kept at room temperature for two nights after pouring. Plates that were not used immediately after this drying period were then refrigerated and used at a later point. Bacteria strains eWM43 and eWM44 were grown overnight from single colonies in 2xYT with ampicillin (100 *μ*g*/*ml), diluted 10^5^-fold in 2xYT with ampicillin and mixed in a ratio of eWM44:eWM43=3:1. For the control experiments without the resistant strain, the dilution of eWM44 was substituted by 2xYT with ampicillin. Four technical replicates of the mixture and two technical replicates of the control (5 *μ*l each) were spotted onto the plate and incubated at 37°C. After roughly 18 hours, phage T7 was added using a pipette tip which was dipped into an aliquot of phage stock. The plate was sealed with Parafilm and individual colonies were continuously imaged using transmitted brightfield and fluorescence every 20 min for 26 hours using a stereomicroscope (Zeiss SteREO Lumar.V12). To provide a constant temperature environment, imaging was carried out in a custom-build enclosure kept at 37°C. During the imaging time course, Parafilm broke, impairing the sealing, but plates did not dry out significantly. This experiment was done in three biological replicates (and technical replicates thereof). All replicates showed the behavior described in the main text. In addition, as is visible in S1 Video for example, lysis of bacteria with red fluorescent marker is regularly visible in the wake of the primary phage front. This is probably due to phage mutants which can infect *E. coli* missing the gene *waaC*. However, this observation does not influence the interpretation of our experiments since these secondary plaques occur well behind the primary front, which is of interest in this work.

#### Plaque growth around obstacles of defined size and shape

A detailed protocol for printing the bacterial lawn is presented in S1 Protocol. In short, agar patches were prepared by placing a 3.5 *×* 2cm^2^ piece of nitrocellulose membrane (Millipore, 0.8 *μ*m AAWP) onto solid agar, covering it with warm molten agar (2xYT medium with 20 g*/*l agar and 100 *μ*g*/*ml ampicillin) and keeping it at room temperature for two nights. Plates were then refrigerated if they were to be used later. The top layer of agar was cut out together with the supporting membrane and placed onto the CD tray of an inkjet printer (Epson Artisan 50) immediately before the printing process (S1 Fig).

Overnight cultures of *E. coli* were grown at 37°C from single colonies in 2xYT with 100 *μ*g*/*ml ampicillin. 5 ml (strain eWM43) and 15 ml (strain eWM44) of overnight culture were spun down and resuspended in 15 ml 42 % glycerol, which was used to fill pristine refillable ink cartridges. (Similarly, cartridges were filled (or refilled) with 70 % ethanol or deionized water and used for cleaning or flushing the printhead in the course of the printing process.) The TIF viewer IrfanView in conjunction with the Epson printer driver was used to print the pattern (CMYK TIF image) onto agar patches, twice for each of the four obstacle shapes. The resulting eight agar patches were than transferred to square plates (same medium as used for agar patches; see S1 Fig and incubated at 37°C for roughly 5 hours. Additional steps allowed us to gauge the sterility and quality of the printing process (see detailed protocol).

At this point, the outline of the obstacle was visible by eye and a linear stretch of phage was inoculated close to the obstacle region using a strip of nitrocellulose membrane (Millipore, 0.8 *μ*m AAWP) soaked with solution of phage stock. (We ensured that no drops were visible on the membrane by stripping the membrane on a sterile surface and/or waiting till the liquid evaporated.) The plate was sealed using Parafilm and the individual regions were imaged using a stereomicroscope (Zeiss SteREO Lumar.V12) inside an custom-built incubator box at 37°C every 20 minutes for at least 24 hours. During the incubation, the Parafilm broke regularly, but the agar surface was not visibly impaired by cracks. Experiments were repeated as biological replicates on three different days with two technical replicates per obstacle shape as noted above (with one exception where placement of the phage-soaked membrane failed). Figs. 3C,D of the main text show the front position and front shape for the replicates of one of the four different obstacle shapes, Figs. 4A,B display the front shape and kink size for all replicates considered. Similar to the experiments of plaque growth in randomly structured heterogeneous environments, we regularly observe lysis of bacteria missing the gene *waaC* (red fluorescence marker). This phenomenon occurs in the wake of the primary front, which is of interest here, and therefore does not affect the interpretation and analysis of our experiments.

#### Spatial expansion of *E. coli* around regions with poor nutrient conditions

Plates with a region of lower nutrient concentrations for *E. coli* were prepared as sketched in S11 Fig: Nitrocellulose membranes (Millipore, 0.8 *μ*m AAWP) with small impermeable regions were created using the Sylgard 184 Silicone Elastomer Kit. A drop of base mixed with curing agent was placed onto the membrane and subsequently cured. (Membranes can be sterilized afterwards using UV light.) Membranes were placed between two layers of medium (2xYT with agar, see below). To that end, 25 ml medium were pipetted into empty standard plates (diameter 8.5 cm). The prepared membrane was placed onto the solidified agar and covered with 2 ml of molten agar and distributed as uniformly as possible. Plates were used after two nights at room temperature.

Bacteria (strains eWM282, eWM284, eWM40) were grown overnight from a lawn or from single colonies in 2xYT with 100 *μ*g*/*ml ampicillin. Next day, overnight cultures were directly mixed or mixed after tenfold dilution in water at a ratio of 1:1:1. A linear inoculation was achieved by soaking a small piece of membrane with a linear edge in the bacterial solution which was then placed onto the agar surface. The plates were incubated at 37°C in a closed container which also contained a beaker filled with water to increase humidity. Plates were imaged regularly using a stereomicroscope (Zeiss SteREO Lumar.V12) in the three fluorescent channels corresponding to the fluorescent proteins used to mark the cells. For analysis, we only considered those experiments where the *E. coli* colony did not completely cover the region with poorer growth condition. The experiments satisfying this criterion were from three different biological replicates, had an unfavorable region with a diameter of between 2.0 and 3.5 mm and the agar concentration was 10, 12.5 or 20 g*/*l and ampicillin was absent or at a concentration of 100 *μ*g*/*ml. All displayed the boost in allele abundance behind the obstacle, sectors meeting just behind the obstacle, and a straight sector boundary behind the obstacle.

### Image Analysis of Bacteriophage T7 Expansions

A semi-automated image analysis pipeline was used to extract front shapes, front positions, and indent sizes (such as in Figs. 3C,D and Figs. 4A,B) from the fluorescence time-lapse information. First, the channel detecting YFP fluorescence was used to define a front right after the plaque boundary got established and the channel detecting mCherry was used to identify the three upper corners of the obstacle. This information was used to define a coordinate system with the the obstacle’s center at the origin and the front extending in *y*-direction (referred to as ‘upper region’ in the following, e.g., Fig. 3B) and the image was cropped (5.3 mm in direction of front movement, 0.7 mm in direction opposite to front movement, and 3 mm to either side of the obstacle). After normalization using the upper, uninfected region, the difference between two consecutive frames (YFP channel) was used to identify the front. In the difference image the extending front manifests itself as a bright region whose upper boundary was identified using thresholding. The algorithm was tested manually since the front is easily detectable by eye, although the decay in fluorescence extends to about 1 mm (S2 Fig). A few frames were excluded from the analysis due to jumps in the front which could be detected automatically using a threshold in local slope of front shape. Finally, front position was determined from the curve of the front close to the boundaries of the cropped region. Indent size was derived as the distance between the most lagging part of the front and front position (after the kink has formed, i.e., curve of front was defined around the axis of bilateral symmetry). The corners of the obstacles identified were also used to identify the size of obstacles, which were slightly smaller than in the printing template. When displaying data for individual obstacles the median of all the obstacles included in the analysis was used; for collapse plots, the size of each single obstacle was used to rescale data. For analysis, front detection was limited to frames obtained within 22 h even if the experiment lasted longer. After this time front detection becomes more challenging most likely due to the bacterial lawn transitioning into stationary phase.

### Numerical Methods & Simulations

#### Generalized Fisher-Kolmogoroff equation - model and parameter choices

To test the generality of the experimentally observed front perturbations and to investigate the lag between the constant speed model prediction and the observed plaque boundary, a reaction-diffusion system was employed, specifically a generalized Fisher-Kolmogoroff-Petrovsky-Piscounoff equation (FKPP equation) [36–39] as outlined in the main text. The FKPP equation was chosen despite the existence of more complex microscopic models for phage spread [31–35] since it is readily parametrizable. As outlined in the main text, two parameters of the FKPP equation,

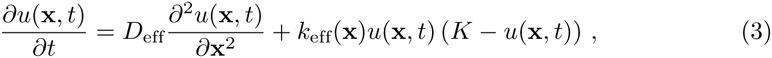

the effective diffusion coefficient *D*_eff_, and the effective growth rate *k*_eff_(**x**) = *k*_0_ specify the front speed via 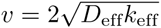 [39] in the case of a homogeneous environment (in our case a homogeneous lawn of susceptible bacteria) and a planar population front. (By rescaling the concentration field *u*(**x***,t*), we can set *K* = 1 without loss of generality.) Conversely, with the front speed determined experimentally, it suffices to know either *D*_eff_ or *k*_eff_ to fully parameterize the model for a homogeneous environment and a planar front. We therefore estimated *D*_eff_ and used front speed far away from the obstacle (*v ≈* 0.2mm*/*h, Fig. 3C) to specify *k*_eff_(**x**) = *k*_0_ outside the obstacle and set *k*_eff_(**x**) = 0 inside the obstacle.

The diffusion coefficient of the phage has not been measured directly for the experimental conditions we employed. Depending on whether the diffusion predominantly occurs in the agar, in the bacterial lawn or in a layer of liquid, the diffusion coefficient is determined by the mesh size of the agar, the density of bacterial cells in the lawn and the humidity. We estimated the diffusion coefficient as follows: First, the diffusion coefficient of phage P22 has been used as a proxy for T7 diffusion [32]; in 10 g*/*l agar *D*_P22_ *≈* 4 *·* 10^*-*8^ cm^2^*/*s = 0.0144 mm^2^*/*h [40]. Second, as an upper bound, we can estimate the diffusion coefficient in water with the phage represented by a sphere with a diameter of 60 nm and a viscosity of 0.692 mPa s for water at 37°C (source: wolframalpha.com) which results in a diffusion coefficient of *D*_SE_ = 0.04 mm^2^*/*h, about a factor three larger than *D*_P22_. Most likely, however, the true diffusion coefficient is smaller than *D*_P22_ since the agar concentration in our experiments is higher (20 g*/*l) and bacteria will restrict diffusion to interstitial regions (see discussion in Ref. [32]). We therefore also considered a diffusion coefficient three times smaller than for bacteriophage P22 in agar as a third case. In all three cases, the front lags behind the prediction of the model of constant speed, see S6 Fig. As expected, for a smaller diffusion coefficient, the lag becomes smaller since the obstacle becomes larger relative to the front width 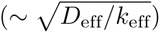 and the constant speed model becomes more accurate. An even smaller value of *D*_eff_ would therefore suggest even better agreement with the prediction of the model of constant speed. Since for our experiments, the value for *D*_eff_ is not known, we chose *D*_eff_ *≈* 4 *·* 10^*-*_8_^ cm^2^*/*s = 0.0144 mm^2^*/*h and therefore *k*_0_ = 0.7 */*h throughout this work. Note that, although the bacterial lawn is changing throughout the experiment (bacteria are presumably not in stationary phase yet), the front speed is roughly constant (Fig. 3C) indicating that small changes in parameters only weakly affect the front dynamics, justifying our coarse-grained approach with these parameter estimates.

Finally, let us note qualitative differences between the coarse-grained model used here and the model of Yin and McCaskill (or rather its extension to heterogeneous environments) [32]: (i) The heterogeneous bacterial lawn enters as a location-dependent parameter of an effective growth rate rather than a heterogeneous initial condition of bacterial density. (ii) While in the coarse-grained model a reduction in density below the carrying capacity due to outward-migration induces growth (Eq. 3), this is not the case for the phage system where growth is not possible in the wake of the front.

#### Generalized Fisher-Kolmogoroff equation - numerical solution

The generalized FKPP equation with a position-dependent growth rate representing obstacles was solved on a square lattice, with periodic boundary conditions along the directions perpendicular to the front on either side of the obstacle and fixed boundary conditions of 1 and 0 well behind and well in front of the obstacle, respectively. The lattice spacing was chosen to be 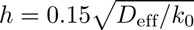. The diffusion operator was discretized using a nine-point stencil (with lattice along *x* and *y* directions: Δ*u*(*x, y*) *≈* [*-*20*u*(*x, y*)+(*u*(*x* + *h, y* + *h*)+ *u*(*x* + *h, y - h*)+ *u*(*x - h, y* + *h*)+ *u*(*x - h, y - h*)) + 4(*u*(*x, y* + *h*)+ *u*(*x, y - h*)+ *u*(*x* + *h, y*)+ *u*(*x - h, y*))]*/*(6*h*^2^).

As initial condition, a linear front profile was chosen, which subsequently developed into the profile for a Fisher wave of width 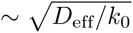 before encountering the obstacle (S4 Video, S13 Fig). The system of ordinary differential equations was solved using the solver ode113 in MATLAB. The front was defined by a population density threshold of *u*_thresh_ = 0.05 (outside the obstacle) with the goal of capturing the edge of the front in a robust manner. (In parallel, we determined the front using *u*_thresh_ = 0._5_. In all cases considered, the front determined using this criterion also showed a lag with respect to the constant speed model, but subsequent analysis was limited to *u*_thresh_ = 0.05.) Front position was defined as the position of the front at the boundaries of the lattice, i.e., far away from the obstacle on either side in the *x* coordinate. To study the effect of a rather irregular obstacle on the shape of the front, the FKPP equation was solved for a mirror-symmetric, non-convex obstacle (S10 Fig). To mimic the obstacles considered in the experiments, the FKPP equation was solved for rhombus-shaped obstacles with (*w* = 0.9mm*,h* = 0.9 mm), (*w* = 0.9mm*,h* = 1.2 mm), (*w* = 1.2mm*,h* = 0.9 mm), and (*w* = 1.2mm*,h* = 1.2 mm). For the obstacle with (*w* = 1.2mm*,h* = 1.2 mm) the lattice spacing, the accuracy requirement of the algorithm, the distance of the boundaries from the obstacle center, and the time until the population wave reaches the obstacle had no influence on front shape.

#### Stochastic simulation of expansion on a hexagonal lattice of demes

To study the effect of the obstacle on allele abundance in the presence of genetic drift, we employed a stochastic model of population growth on a hexagonal lattice, specifically a set of demes with zero or one organisms per deme and stochastic growth onto neighboring sites (see, e.g., [5, 45]). A lattice model where the shape of the front has no undulations simplifies analytical calculations [5], but is not applicable here since the shape of the front is a priori unknown. We therefore generalized the model of unconstrained growth in two dimensions as described by Korolev et al. (Fig. 2A in Ref. [5]) to more than two alleles, see S12 Fig for a detailed description. The obstacle was realized by withdrawing the ability to serve as empty or inhabitable lattice sites for a set of lattice sites (marked in gray in Fig. 5C and S12 Fig). The same holds for the boundaries perpendicular to the overall front movement. (We did not choose periodic boundary conditions to simplify the illustration of the ancestry in Fig. 5C.) The system width was chosen to be approximately 600 times and the obstacle’s radius to be about approximately 80 times the distance between adjacent lattice sites. The obstacle center was placed one third of the system width ahead from the originally populated lattice sites and in the middle between the two boundaries to the left and right. One occupied row on the hexagonal lattice with all unique alleles (i.e., colors) served as initial condition. In the course of the simulation the front with these rules roughens somewhat, an aspect which is not of central interest here. The ancestry of all occupied lattice sites with at least one free neighboring lattice sites was determined by keeping track of the history of the simulation at different points in time (Fig. 5C).

We set up ten instances of the simulation and interpreted the results qualitatively. The following features were clearly visible in frames which included the ancestry for all instances or for the majority of instances: dynamics of the front as described throughout this paper, the loss of alleles encountering the obstacle head-on, an increased abundance in the shade of the obstacle of alleles grazing the obstacles, two sectors meeting at top of the obstacle, the lineages from individuals at the curved front originating from around the point of maximum width, and the ancestry lines not passing through the point at the top of the obstacle at larger times.

## Supporting Information

### S1 Video

#### Time-lapse movie of phage spread in heterogeneous environment

(see Fig. 2A for snapshots, see Figs. 1C,D for snapshots of fluorescence channels only). As Fig. 1B illustrates, yellow regions represent a bacterial lawn of *E. coli* cells susceptible to infection with bacteriophage T7 while red regions represent resistant *E. coli* cells.

### S2 Video

#### Control experiment for phage spread in heterogeneous environment

Like S1 Video, but without patches of resistant cells. The shape of the plaque is much more regular.

### S3 Video

#### Time-lapse movie of plaque extension on a printed lawn

(dark region) of susceptible bacteria (yellow) in the presence of an obstacle, a region of resistant bacteria (red). (For snapshots see Fig. 3B.)

### S4 Video

#### Visualization of the numerical solution of the reaction-diffusion model for an rhombus-shaped obstacle

with *w* = 1.2mm and *h* = 0.9 mm (scale bar: 1 mm). The obstacle, region of growth rate *k*_eff_(**x**) = 0, is indicated in red, population density *u* is indicated in green. On the right, the profile of the front on the left and right boundary of the lattice is shown. Scale bar represents 1 mm. (For snapshots see Fig. 4E.)

### S5 Video

#### Visualization of the numerical solution of the reaction-diffusion model for an obstacle with a more complex shape

For details, see caption of S4 Video, for snapshots see S10 Fig.

### S6 Video

#### Expansion of an *E. coli* colony with three different strains with three different fluorescent markers around a region with poorer growth conditions

over the course of 13 days. (For snapshots see Fig. 5B.)

### S7 Video

#### Visualization of a stochastic simulation of a population with neutral alleles in the presence of an obstacle

The obstacle is denoted by a gray circle, initially about 600 alleles are present. Black lines in the last frame (and the snapshots in Fig. 5C) represent lineages of lattice sites at the front.

### S1 Appendix

The constant speed model: Additional results, analogy to geometrical optics, and limitations

See below.

### S1 Protocol

Detailed protocol including general considerations for printing the bacterial lawn

See below.

**Fig S1.**
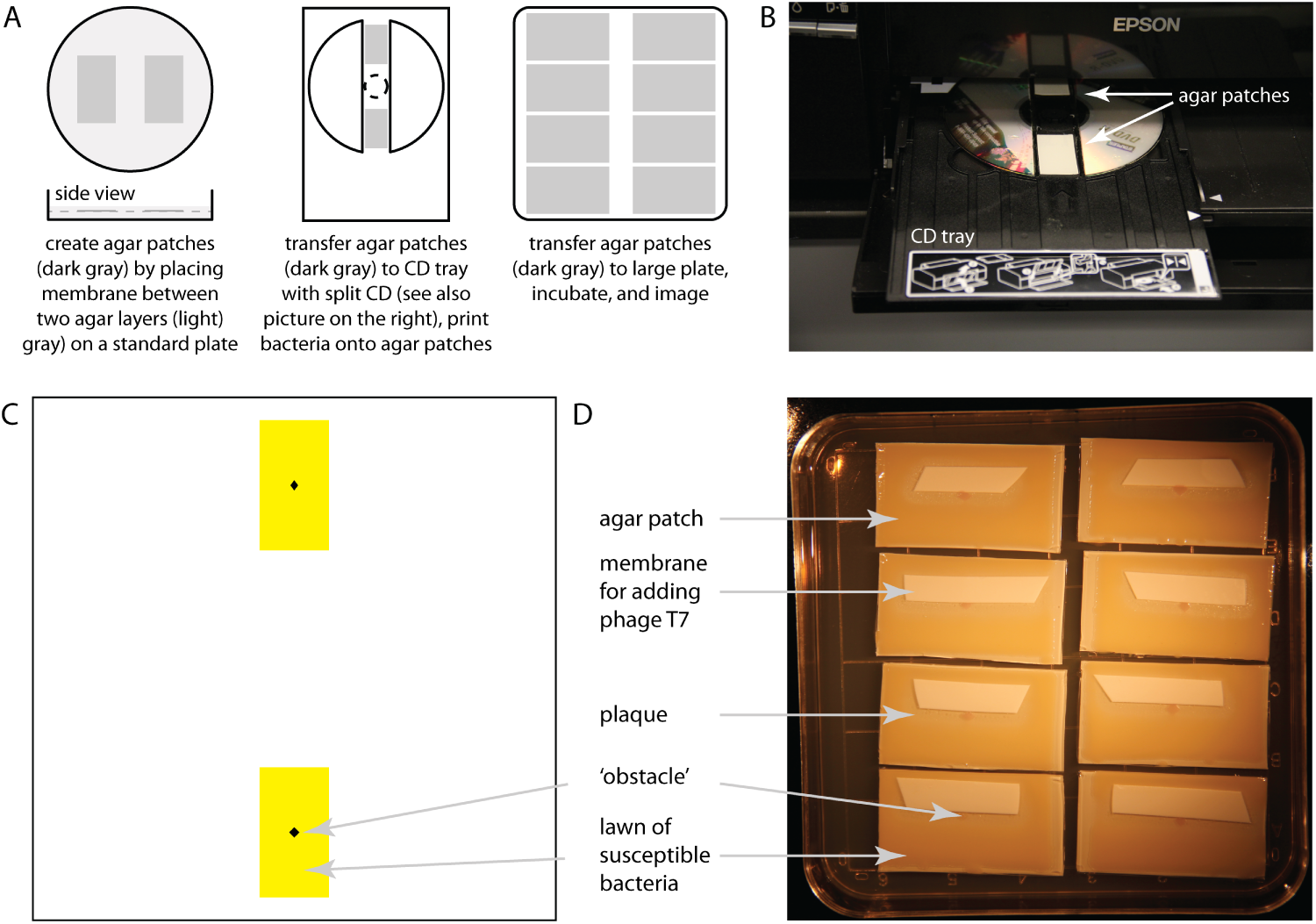
Illustration of the printing protocol. **(A)** First, agar patches are created by placing a membrane (cut to 3.5 *×* 2mm^2^) between two layers of agar. The membrane with the top layer of agar is cut out and placed onto the printer’s CD tray. A split CD is used to define position and ensure proper functioning of the printer. After printing of the bacterial solution, the agar patches are transferred to a larger agar plate which then is incubated. **(B)** Photograph of the CD tray with two agar patches loaded into the printer (Epson Artisan 50). **(C)** Image of the pattern used to print bacteria onto the agar patch. Susceptible bacteria are found in yellow cartridge, resistant bacteria in black cartridge which after printing leads to the pattern described in Fig. 3A. **(D)** Picture of the plate with experiments on plaque growth around single obstacles, after incubation and imaging.

**Fig S2.**
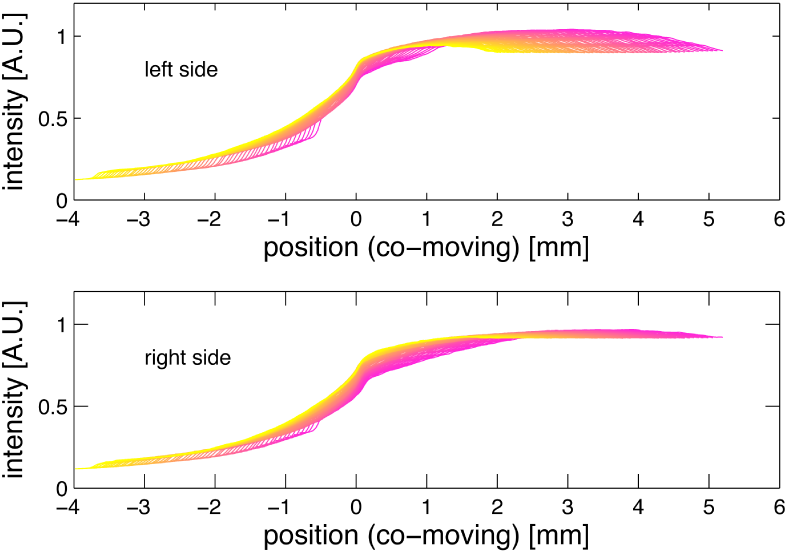
Profile of fluorescence intensity in the channel representing the cells susceptible to bacteriophage T7 infection. 3 mm to left and right of the center of the rhombus-shaped obstacle in the co-moving frame (where the front position *y* is to be at 0) for the experiment displayed in Fig. 3B and S3 Video. Different colors represent different time points where yellow corresponds to late times after the start of the experiment. Different front profiles do not collapse onto each other perfectly (and do so less well in other experiments), but the small differences are not expected to influence our results on front shape since algorithmic front detection was checked manually.

**Fig S3.**
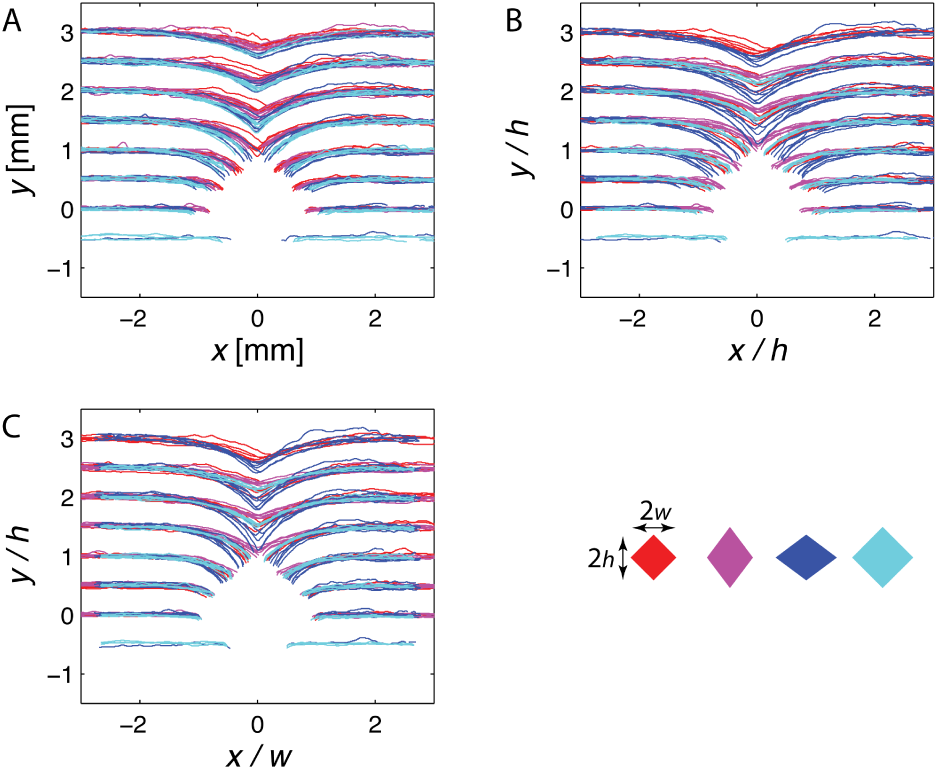
Experimentally determined front shapes from biological and technical replicates for different obstacle shapes as indicated by color. Axes are not scaled or scaled by half the obstacle width, *w*, or half the obstacle height, *h*, as indicated (see also Fig. 4A in main text). Collapse is best when all length are measured in units of *w* (Fig. 4A in main text) as predicted by model of constant speed. The collapse of front shapes for obstacles of same width (but not same height) in (A) illustrates that width, not height, of the obstacle determines front shape for rhombus-shaped obstacles.

**Fig S4.**
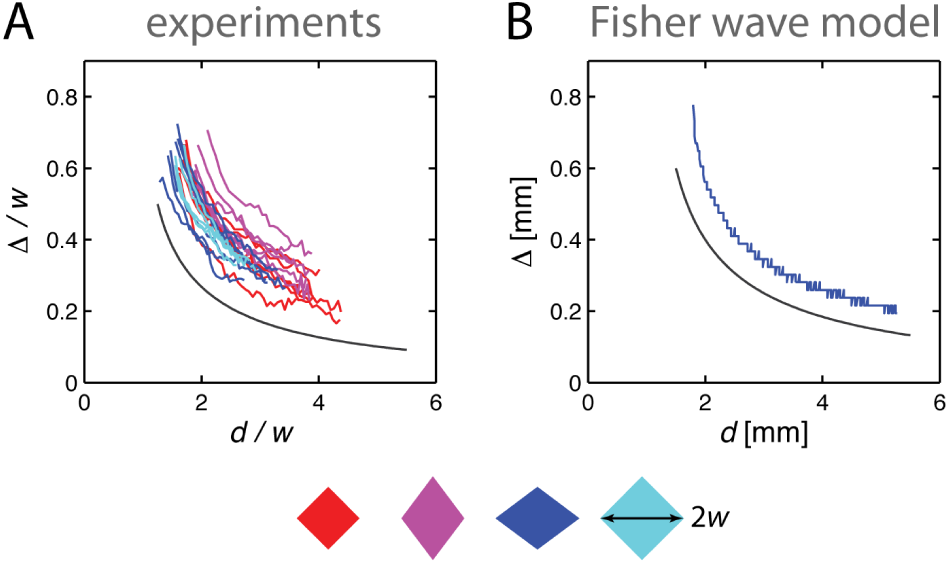
Indent size Δ as a function of front position. *d* for both experiments and the numerical solution of the reaction-diffusion model for different types of obstacles specified below by color. The experimental data are scaled by half the obstacle width, *w*, compare Figs. 4B and D of the main text.

**Fig S5.**
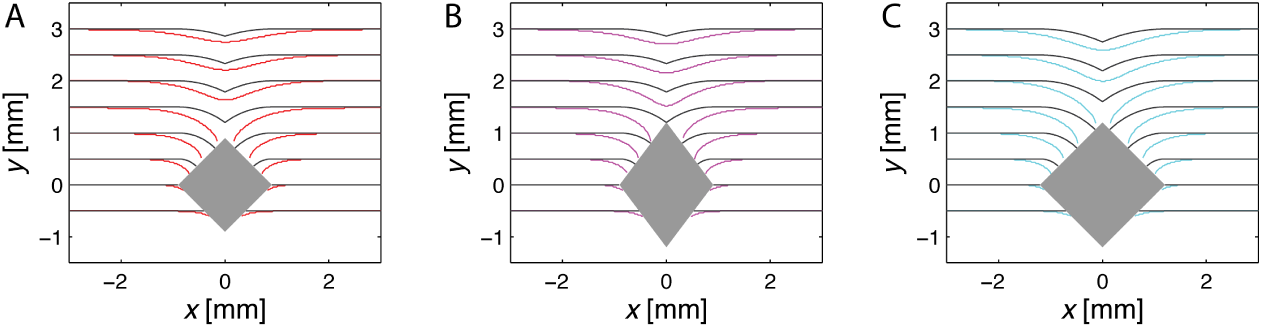
Front shape in the reaction-diffusion model for different obstacle shapes. (colored line) and the model of constant speed (black line), see Figs. 4C,D in the main text. In all cases, the front in the reaction-diffusion model lags behind the front in the model of constant speed.

**Fig S6.**
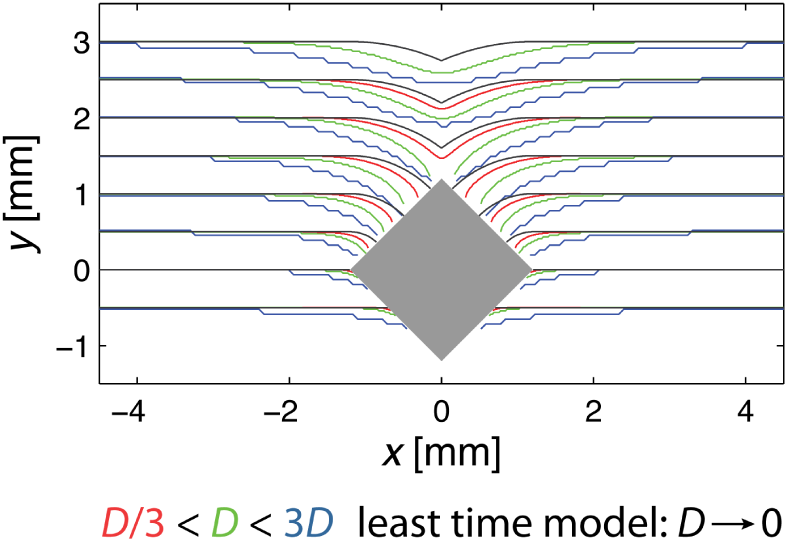
Front shape in reaction-diffusion model with different effective diffusion coefficients. *D*_eff_ (colored lines) and model of constant speed (black line). (Choice of *D*_eff_ determines effective growth rate *k*_0_ via 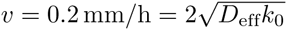 as explained in the main text). Green: *D*_eff_ = *D* = 0.0144 mm^2^*/*h, red: *D*_eff_ = *D/*3, blue: *D*_eff_ = 3*D*, see Materials & Methods for justification of the choices for *D*_eff_. In all cases, the front in reaction-diffusion model lags the front predicted by the model of constant speed. The reduction of the lag with decreasing *D*_eff_ is consistent with a decrease in *ξ* which characterizes the limits of the model of constant front speed.

**Fig S7.**
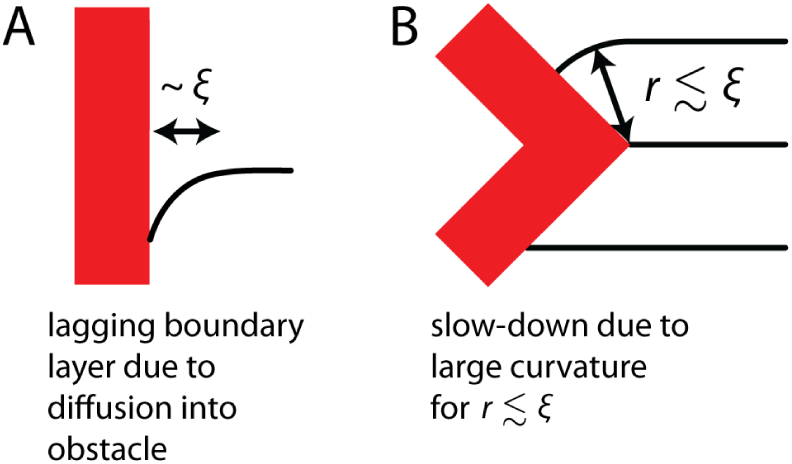
Mechanisms for a lag of the front. **(A)** Diffusion of the reproducing phage into the obstacle leads to a lagging front due to a boundary layer of width 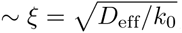, where *D* is the diffusion coefficient and *k* the local growth rate. **(B)** A (rapid) change in the slope of an obstacle boundary can induce a lag while the radius *r* of the circular segments of the population front is smaller than the characteristic length scale *ξ*, *r* ≤ *ξ*. See text for details.

**Fig S8.**
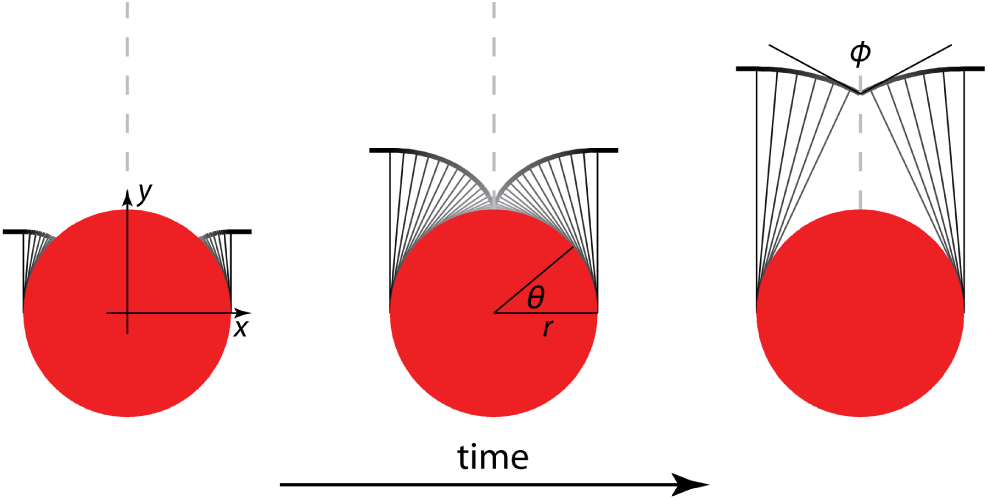
Geometric construction of the front behind a circular obstacle of radius *r* using the constant speed model, (from left to right) before the kink forms, when the kink forms from a cusp with infinite vertical slope and as the kink heals. At any given time, the front in the shadow of the obstacle is composed of infinitely many circular segments with centers at the boundary of the circle. Since the upper boundary of the obstacle is parallel to the original front, the opening angle *Φ* of the kink vanishes at the moment the kink forms, resulting in two locally parallel population fronts. (The kink therefore is a cusp in this case.) As the kink heals, circular inflation still occurs locally, but the arc length of the perturbed front is reduced. At large times during the healing phase fewer and fewer circular segments contribute to the front: Finally, only those with centers close to the point of maximum width are relevant. In this long time, large distance limit, the detailed shape of the obstacle drops out. In addition, the coordinate system and the parameter *θ* used to describe the circle are indicated.

**Fig S9.**
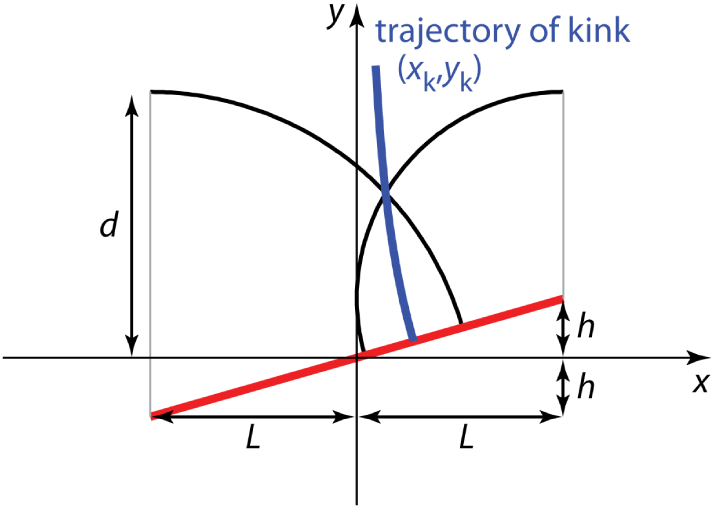
Illustration of the front shape for a tilted, very thin obstacle, which is indicated by the red line. The projected width of the obstacle is 2*L*, the obstacle is tilted by an angle arctan(*h/L*). Black segments of circles centered on the edges of the obstacle indicate front shape at a given point during the healing process. The kink forms off-center, but its tip approaches a line normal to the unperturbed population front that bisects the projected obstacle (curved blue line).

**Fig S10.**
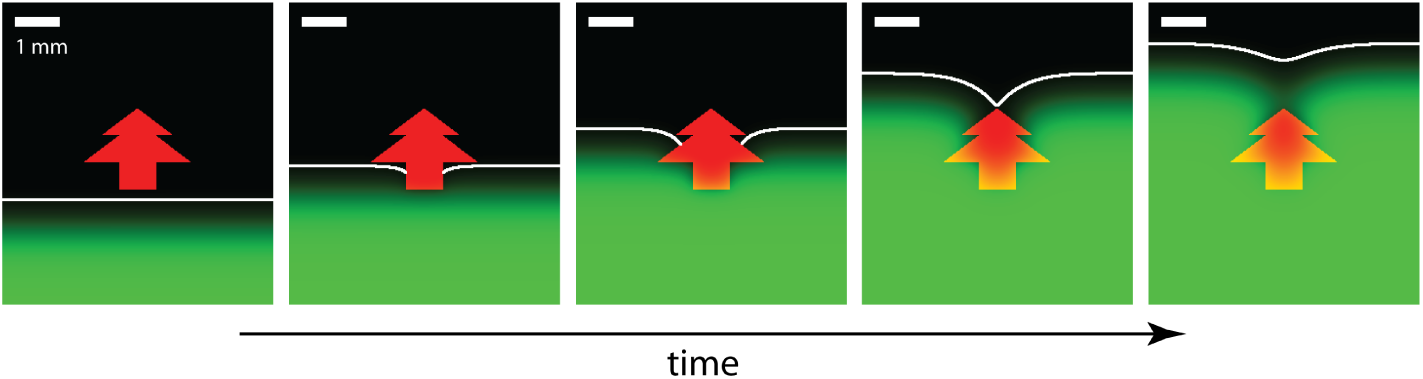
Snapshots of the numerical solution of the reaction-diffusion model with a region of no growth with complex shape. The region of no growth is indicated in red. Population density is indicated in green, the inferred front of the traveling population wave is marked white. See S5 Video for all frames and compare to Fig. 4E for a rhombus-shaped obstacle.

**Fig S11.**
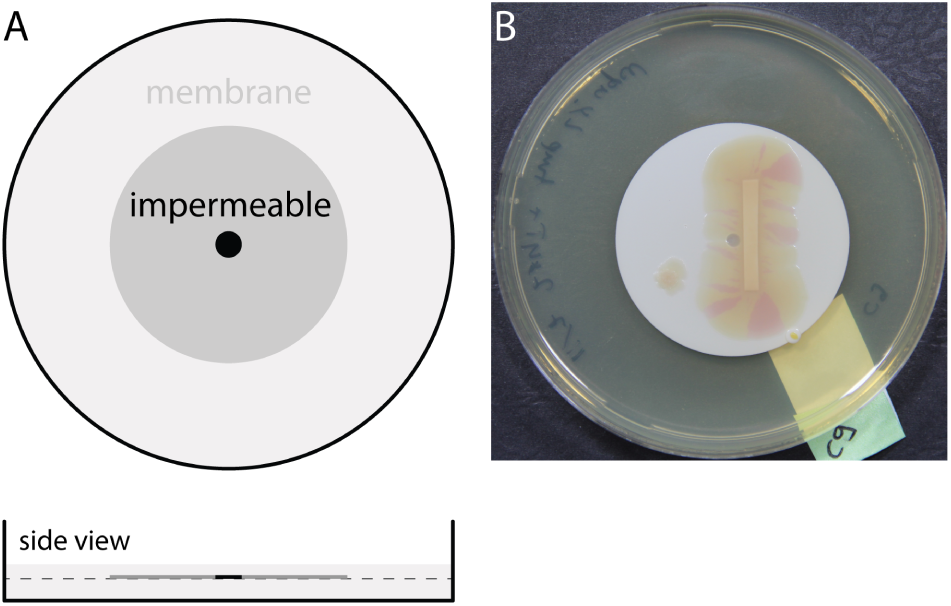
Experimental setup for bacterial expansions in the presence of a region with poorer growth conditions. **(A)** A part of a membrane is first made impermeable (see Materials & Methods). The membrane is then placed on top of an agar layer and covered by a thin layer of agar. **(B)** Picture of plate after bacterial expansion took place.

**Fig S12.**
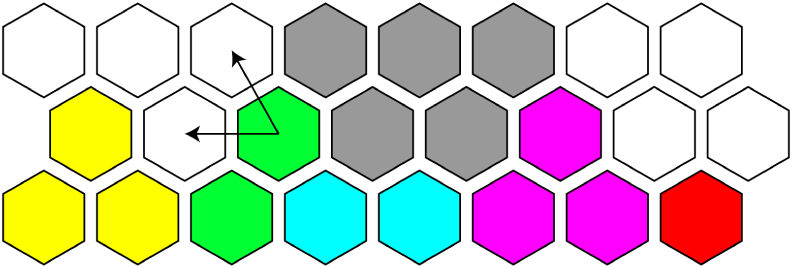
Illustration of the algorithm for the stochastic simulation with one organism per deme. (an extension of an Eden model variant including different genotypes, see also Fig. 2a of Ref. [5]). On a hexagonal lattice, each lattice site can be occupied by one organism with a given allele, represented by the lattice site being assigned a given color. The organism (or the lattice site) can reproduce by converting a neighboring, empty lattice site into a site of the same color. In this process, a site with at least one empty neighboring site is chosen at random (here: a green one) and randomly converts one of the empty neighboring sites (two possibilities, indicated by arrows). The gray sites representing the obstacle cannot be occupied and are regarded as filled non-reproducing sites in the simulation.

**Fig S13.**
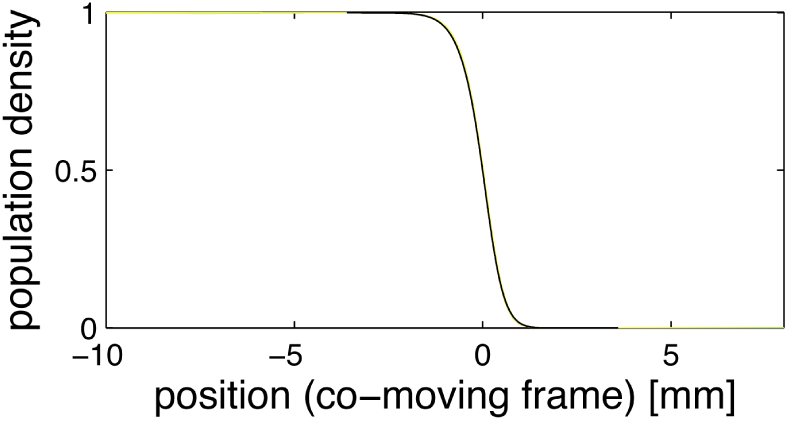
Shape of the population profile far away from the obstacle for the reaction-diffusion model. Profile is determined at the boundary of the lattice for the numerical solution displayed in S4 Video after the front has encountered the obstacle. The black line indicates the approximation *u*(*z*) *≈* (1 + *e*^*z/c*^)^*-*1^ +1*/*4 *· e*^*z/c*^(1 + *e*^*z/c*^)^*-*2^ ln(4*e*^*z/c*^(1 + *e*^*z/c*^)^*-*2^) with 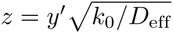, where *y’* is the position in the comoving frame, *k*_0_ and *D*_eff_ are the growth rate and diffusion coefficient as specified in Materials & Methods, and *c* = 2 is the dimensionless front speed; see Ref. [39] for more details.

## Acknowledgments

We thank Charles C. Richardson and Seung-Joo Lee for an aliquot of wild-type T7 and discussions; we are grateful to John Chuang, Daniel Cohen, Clément Vulin, and Christoph Riedl for helpful suggestions.

## S1 Appendix: The Constant Speed Model

## 1 Constant Speed Model for Obstacles of Different Shape

**1.1 Rhombus-shaped obstacle**

Assume a rhombus with width 2*w* and length 2*h* with its width parallel to a population front encountering this obstacle as sketched in Fig. 2G. The position of the population front at a given time is characterized by the distance *d* traveled normal to the front (Fig. 2E). The constant speed model predicts that the front remains planar until the point of maximum width is reached (conveniently defined as *d* = 0, see main text and Fig. 2E). If the speed *v* of the front remains constant over the relevant time interval, we have *d* = *vt*. More generally, 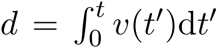. Note that the prediction of the front shape is independent of *v*(*t*), but we chose the term ‘constant speed model’ for clarity. The curved part of the front, in the shadow of the obstacle, is given by the arcs of two circles with radius *d*. A kink forms at 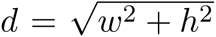 and is characterized by opening angle *Φ* (see S8 Fig for the analogous opening angle *Φ* for a circular obstacle) and indent size Δ. Both are independent of *h* (as is front shape, see Fig. 2G) and are given by:

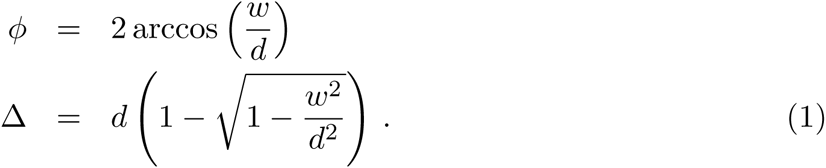

For *d≫ w* we obtain

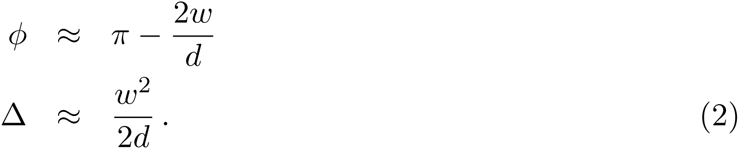

**1.2 Other convex obstacles with bilateral symmetry**

Aligned rhombus-shaped obstacles are simple in the sense that the constant speed model predicts initiation of two radial population waves simultaneously at *d* = 0, when the traveling wave passes the point of maximum width. This simplicity also holds true for rectangular obstacles (aligned parallel to the population front), where radial population waves are initiated when the front has just finished grazing the sides of the obstacle. For more complex obstacles, multiple radial population waves can be initiated. Let us first consider general convex obstacles with bilateral symmetry before quantifying the considerations for circular and elliptical obstacles below (see S8 Fig).

According to the constant speed model, the front of the population wave at distance *d* is determined by an ensemble of paths of length *d* all hugging the boundary before continuing tangentially. There is thus a successive initiation of circular population waves along the boundary of the obstacle. Hence, the tangents to the front and the boundary of the obstacle form a 90° angle after the front has passed the point of maximum width as shown in the left-most part of S8 Fig.

For long times (and large distances downstream from the obstacle) the front is completely determined by segments originating around the region of maximum width (S8 Fig). The front in this limit is thus only determined by the shape of the obstacle near the point of maximum width, and ultimately only by the width of the obstacle 2*w*. Hence, asymptotic results such as Eqs. 2 hold independent of the detailed obstacle shape.

**1.3 Circular obstacle**

The above considerations for convex shapes with bilateral symmetry are conveniently illustrated with a circular obstacle. (The more general case of elliptical obstacles will be treated later.) For a circular obstacle (radius *r*) it is convenient to use the polar angle *θ* to parametrize the boundary (S8 Fig). From the constant speed model sketched above, we find the coordinates for the right part of the front parametrically as function of *θ* at fixed 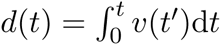 as

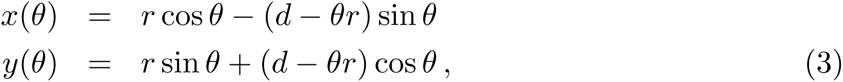

where *x*(*θ*) and *y*(*θ*) are given relative to the center of the obstacle.

The results hold for all positive distances *d* relative to the midline of the circle parallel to the initial front, but the maximum value of 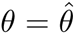 allowed in Eq. 3 depends on how far the wave has traveled:

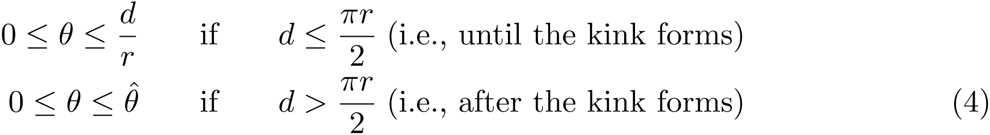

where 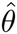 is determined by the condition 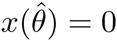

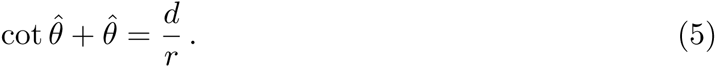

The kink shape at its birth follows from expanding *x* and *y* around *θ* = *π/*2 and eliminating *θ*. The cusp shape that marks the birth of the kink is given by

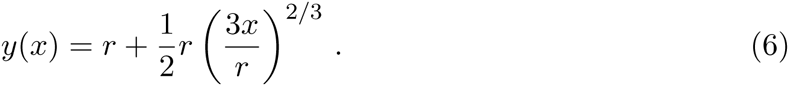

Note that the initial opening angle *Φ* vanishes in this case, i.e., *y*(*x*) has an infinite slope.

The subsequent healing of the kink can be characterized similar to rhombus-shaped obstacles by the indent size Δ and opening angle *Φ*:

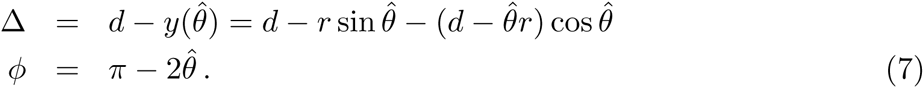

The implicit Eq. 5 for 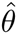 cannot be solved in closed form. However, one can consider the case of *d/r ≫* 1 (i.e., the front has already traveled several obstacle lengths downstream) which implies 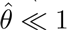. We make a polynomial ansatz for 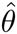 as

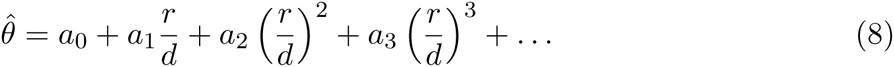

Expanding the inverse Eq. 5 and comparing coefficients on both sides of the equation we find

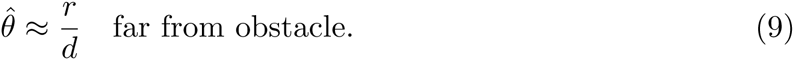

Employing the polynomial representation of 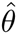 and keeping the lowest terms depending on *r/d* we obtain for opening angle *Φ* and indent size Δ:

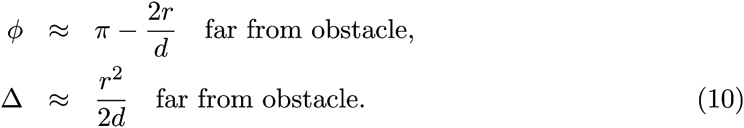

Note that upon identifying *r* with *w* (half the maximum width of the obstacle), the opening angle and indent size of the kink agree with our earlier result, Eq. 2, for rhombus-shaped obstacles, consistent with universal large distance behavior for the kink.

**1.4 Elliptical obstacle**

The shape of the front encountering an obstacle shaped as an ellipse (oriented with one axis parallel to the unperturbed front) can be derived similarly. Assume here that the front encounters the ellipse parallel to the major axis of length 2*a* and with front direction parallel to the axis of length 2*b* with *a > b*. A parameter *τ*, 0 *≤ τ ≤* 2*π*, can be used to parametrize both the shape of the ellipse and the curve of the impinging front. Upon taking the center of the ellipse as the origin the boundary of the ellipse is given by *x*_el_(*τ*) = *a* cos *τ* and *y*_el_(*τ*) = *b* sin *τ*. To compute the shape of the front we need the tangent to the ellipse and the arc length of the ellipse’s boundary where the path is hugging the obstacle, both of which can be derived from the parametric form. In analogy to the circular case we find for the coordinates *x*(*τ*) and *y*(*τ*) of the right part of the front:

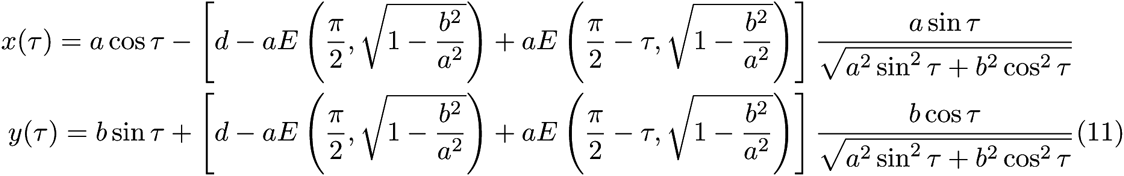

where we used the incomplete elliptic integral of the second kind [1],

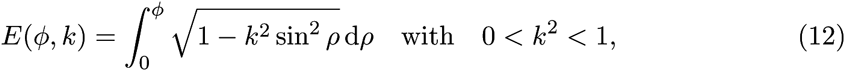

to simplify the expression. Note that the parameter *τ* parametrizing the ellipse is similar but not identical to the polar angle 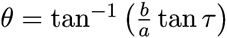 measured from the ellipse center.

The kink forms when the wave has traveled a distance 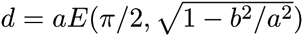, along one quarter of the ellipse perimeter. Before the kink forms, the upper limit of *τ* is given by the condition that the expression in square brackets in Eqs. 11 is larger or equal to 0 while after formation of the kink the upper limit of *τ* is given by *x*(*τ*) *≥* 0.

To determine the shape of the kink at the moment it forms as a cusp one can set 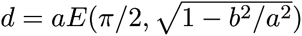 and *τ* = *π/*2 *-∊*, expand for small *∊*, and express *y* as a function of *x* resulting in:

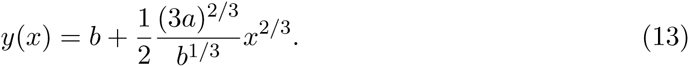

Again, note the vanishing opening angle and the characteristic power law *y*(*x*) *- b ∼ x*^2^*/*3.

**1.5 Tilted, convex, symmetric obstacles**

So far, only obstacles with bilateral symmetry relative to the front direction were considered. How do deviations in alignment affect the kink trajectory? To answer this question, we can use the constant speed model as shown in S9 Fig. Let us consider a thin flat obstacle with projected length 2*L* (onto the incoming front) and a vertical offset by 2*h* (resulting in a tilting angle of arctan(*h/L*)). The trajectory of the kink, (*x*_k_*,y*_k_), is given by the intersection of two circles, one with radius *d* + *h* centered at (*-L, -h*) and another with radius *d - h* centered at (*L, h*), where *d* is the distance of the front traveled in *y*-direction relative to a line parallel to the unperturbed front passing through the center of the obstacle. The resulting kink trajectory is

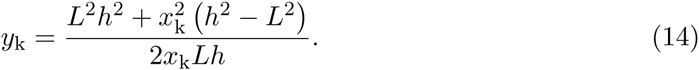

As indicated in S9 Fig, for large distances beyond the obstacle, the kink position slowly approaches the *y*-axis bisecting the projected obstacle, along a hyperbolic curve.

## 2 Analogy to Geometrical Optics

The constant speed model has a strong analogy to geometrical optics. The blue lines in Fig. 2G which mark the path of an imaginary particle at the front can be found by minimizing the path back to the front, which is equivalent to *Fermat’s principle* minimizing the integral over optical density to infer the light path between two points [2, 3].

While the analogy to Fermat’s principle is intuitive from a mathematical perspective, especially for extending predictions to more complex environments, the use of *Huygens’s principle* gives an intuitive explanation for the construction of the wave front (black lines in Fig. 2G). Each point along the front is the source for a radial wave. The envelope of all these waves determines the wave front at a later time which intuitively explains the circular shape of the front in the shade of the obstacle [2]. Note that, of course, there is no interference as described by the Huygens-Fresnel principle [2].

Based on these analogies we also predict that the influence of non-perfect obstacles, which support traveling population waves with smaller speed, can be described by such a minimization procedure. Specifically, we believe that Snell’s law of refraction (which can be derived from either description of geometrical optics) holds for population waves traversing boundaries between habitats in which the populations propagate with different speed.

Last, as explained in the main text, the constant speed model only is a good description of front shape if the size of the obstacle *L* is much larger than the front width parameter *ξ*. The analogous condition for geometrical optics is *λ ≪ L*, where *λ* is the wavelength.

## 3 Origin of the Front’s Lag - Limitations of the Constant Speed Model

S7 Fig illustrates two mechanisms which lead to a lag of the front relative to the constant speed model prediction as outlined in the main text. First, diffusion of phage into the obstacle leads to a local reduction of population density close to the boundary and in consequence to a lagging part of the front (S7 Fig, panel A). This lagging region forms a boundary layer with a width of the order of the diffusion length 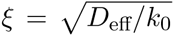. Unless *L ≫ ξ*, the boundary layer is significant and can lead to an apparent lag compared to the prediction of the model of constant speed. As *L/ξ → ∞*, the boundary layer still exists, but is only a minor perturbation of the overall front shape. Although phage diffusion into the obstacle most likely plays a role in our experiments, note that experimentally observed fronts encounter the boundary at an angle more closely resembling 90° (Figs. 3B,4A) than the population front originating from the FKPP equation (Fig. 4C). Although we attribute this difference to the coarse-graining of our model (and differences in front detection), other effects in addition to the boundary layer could contribute to the lag. Further insights might arise from repeating the experiments and numerics using an obstacle with a reflecting boundary.

A second limitation arises because the constant speed model for rhombuses predicts a sudden change in the curvature of the front close to the position of maximum width, right when the circular arc arises (S7 Fig, panel B). Large curvatures can influence front speed of a Fisher population wave. To understand this effect, consider the case of uniform curvature, i.e., a radially expanding population in two dimensions. In radial coordinates and for a radially symmetric population density *u*(*r*) the FKPP equation reads: *∂u*(*r, t*)*/∂t* = *D*_eff_*∂*^2^*u*(*r, t*)*/∂r*^2^ + (*D*_eff_*/r*) *· ∂u*(*r, t*)*/∂r* + *k*_0_*u*(*r, t*)(1 *- u*(*r, t*)). The second term only contributes significantly for radii of order the characteristic length *ξ*, but then leads to a slow-down of the wave [4]. We therefore expect a contribution to a lag of the front whenever circular arcs with initially very small radius are generated. As illustrated in S7 Fig (panel B), for a rhombus this is the case at the point of maximum width. We expect the relative contribution of this effect to front shape to decrease with increase of the obstacle size while keeping obstacle shape constant: For a rhombus, the slow-down due to high curvature occurs when the front grazes the obstacle for small and large obstacles alike, but the overall perturbation of the front is larger for larger obstacles, diminishing the importance of the lag. For obstacles without sharp corners but smooth boundaries, such as the circles and ellipses considered above, the constant speed model also predicts the emergence of circular arcs with very small curvature, in these cases all along the boundary. We therefore also expect a lag due to large curvature of the population front close to the obstacle’s boundary. More work is needed, however, to quantify the effect of this continuous generation of arcs with large curvature along a smooth boundary and its dependence on the the obstacle’s size.

## S1 Protocol: Printing the Bacterial Lawn

In short, bacteria are printed from custom-filled ink cartridges onto agar patches placed onto the CD (Compact Disk) tray of the Epson Artisan 50 printer. Agar patches are created with a supporting membrane underneath. Cartridges are filled with bacterial cells suspended in 42 % glycerol and are used to print bacteria onto the agar patch from a CMYK TIF image prepared with MATLAB. Additional cartridges with deionized water and 70 % ethanol and original Epson cartridges with ink are used to flush the printhead, clean the printhead from bacterial solution and as a test of printhead function. S1 Fig displays the experimental setup and procedures. The following sections provide detailed protocols.

### 1 Choice of printer

For our experiments, an Epson Artisan 50 printer was used, since the protocol described here is based on the approach used in Ref. [1] (using an Epson R280 printer, a very similar model). In general, a printer is needed which (i) is able to print on CDs, (ii) uses piezoelectric material (instead of heat) to generate droplets (arguably more gentle to cells printed), and (iii) is able to print on CDs and therefore has a CD tray on which agar patches can be placed during the printing process.

### 2 Customizing the CD tray

The CD tray of the Epson Artisan 50 printer serves as the supporting substrate on which the agar patches are located during the printing process (S1 Fig). Parts of a cut-up CD, glued into the CD tray, serve as spacers between the CD tray itself and components within the printer. With the position of CD parts and the agar patches carefully chosen, it is possible to minimize contact between the agar patch and the parts inside the printer which move and fixate the CD tray. Note that it might be necessary to cover openings in the CD tray which potentially serve to detect the absence of a CD by the printer driver. Preparation of several trays as support for the agar patches enables quick processing of replicate experiments. Trays were cleaned with detergent, rinsed with water and rubbed with ethanol to minimize contaminations.

### 3 Making agar patches and plates

1. Prepare warm molten agar: liquid 2xYT medium with 20 g*/*l agar and antibiotics as needed.
2. Produce agar patches: Pipette 10 ml of medium into standard plates (diameter 8.5cm, place a 2 *×* 3.5cm^2^ piece of nitrocellulose membrane (Millipore, 0.8 *μ*m AAWP) onto the solidified agar. Pipette 5 ml on top and distribute as well as possible.
3. Supporting plates: Pour 40 ml of medium into square plates (9 cm *×* 9cm).
4. Keep plates in the dark at room temperature for two nights, then refrigerate if not used immediately.

### 4 Mapping from pattern to cartridges

When using a consumer inkjet printer to print a customized liquid, one challenge is to ensure the best possible mapping between colors used in the pattern and the cartridges used to print the pattern. One approach is to print one component at a time to avoid depositing liquid of the wrong type by the printer driver. For example, in an attempt to print a rich yellow, the printer driver might add some liquid from the black cartridge. We did not follow the approach of printing from one cartridge at a time [1] because it vastly slows down the printing protocol and small numbers of resistant bacteria in a region of predominantly susceptible bacteria do not influence the experiment and vice versa. As a practical matter, we found it easiest to design a template as a CMYK TIF image in MATLAB, which was printed using IrfanView and the Epson printer driver using Windows 7.

The Epson Artisan 50 printer prints with six cartridges containing yellow, black, cyan, magenta, light cyan, and light magenta inks. The yellow and black channels and specific driver options were heuristically chosen to minimize ambiguities in the assignment of colors to cartridges: ‘CD/DVD’, ‘Ultra Premium Photo Paper Glossy’, ‘Photo’, ‘A4 (210 x 297 mm)’, ‘Borders’, ‘Portrait’, ‘Fix Red-Eye’ unchecked, ‘High Speed’ unchecked, ‘Edge Smoothing’ unchecked, ‘Print Preview’ unchecked, ‘Black/Grayscale’ unchecked, ‘Color Management’ : ‘ICM’ & ‘Off (No Color Adjustment)’. Margins were adjusted to ensure the pattern was printed where the agar patches were placed, which can easily be tested by printing ink on paper mimicking the agar patches.

In the course of developing the printing assay the software package Gutenprint, providing an alternate printer driver, was also used to print patterns. The settings Gutenprint provides can be used for a non-ambiguous mapping, but this approach was not followed up in the course of this work.

### 5 Filling and refilling cartridges

- Refillable cartridge set T0781-T0786 from InkjetMall (East Topsham, VT) was used for our experiments. Unfortunately, this product is no longer sold, but any refillable cartridge set compatible with the printer of choice should work for the protocol outlined above.
- Cartridges were filled deionized water, 70 % ethanol and bacterial solution as described by the manufacturer. Note that filling the cartridges for the first time might differ from the procedure used to refill the cartridges.
- Cartridges with bacteria were used only once, cartridges with deionized water or ethanol were refilled when necessary and if contaminations could be ruled out.
- The protocol outlined below requires significantly more yellow and black cartridges than cartridges of any other color. Costs can be reduced by swapping the chips between cartridges (if possible), which allows one to use cartridges originally meant for other colors than yellow or black to be used with bacterial solution.

### 6 Printing bacteria onto agar patches

The protocol below reflects the printing assay and cartridges used in this paper, but can easily be adjusted to print two arbitrary bacterial strains.

1. The day before: Make overnight culture of eWM43 and eWM44 from single colony in 17 ml 2xYT with 100 *μ*g*/*ml ampicillin.
2. Spin down 5 ml of eWM43 and 15 ml of eWM44 culture. Resuspend cells in 15 ml of 42 % glycerol by vortexing.
3. Fill empty cartridges with bacterial solution using a syringe by creating a vacuum inside the cartridge which is replaced by the bacterial solution. Remove air vent tabs of cartridges afterwards.
4. With Epson cartridges in printer, perform nozzle check as provided by Epson software. Ensure all nozzles are working properly.
5. Replace cartridges by custom cartridges filled with deionized water. Perform ‘head cleaning’ twice to flush printhead. Perform nozzle check to ensure no ink is left in the printhead.
6. Print one pattern using software IrfanView on agar patches placed on CD tray with printer options as specified above. Transfer agar patches to agar-filled plate and incubate at 37°C. Here, deionized water is printed and incubation of these patches thus serves to test for contaminations.
7. Replace yellow cartridge filled with deionized water by yellow cartridge filled with bacterial solution (eWM43, *E. coli* susceptible to bacteriophage T7 infection). Perform ‘head cleaning’ twice to flush bacterial solution into the printhead.
8. Print one pattern using the same settings as before and transfer agar patches to agar-filled plate. These patches serve as an estimate of how many susceptible cells are printed within the obstacle. After transfer to an agar-filled plate and incubation at 37°C even single cells will occur as colonies which are easily detectable under a microscope. Note, however, that this test only concerns the printer driver and does not consider that printing properties can change when the the black cartridge filled with deionized water is replaced by a black cartridge filled with bacterial solution (next step).
9. Replace black cartridge filled with deionized water by black cartridge filled with bacterial solution (eWM44, *E. coli* resistant to bacteriophage T7). Perform ‘head cleaning’ twice to flush bacterial solution into the printhead.
10. Print different patterns (with two obstacle shapes each) onto patches and transfer agar patches to agar-filled plate. Incubate at 37°C before adding phage and imaging as explained in Materials & Methods.
11. Insert cartridges with 70 % ethanol into printer. Perform ‘head cleaning’ twice to clean printhead.
12. Insert original Epson cartridges into printer. Perform ‘head cleaning’ twice to flush original Epson ink into printhead.

During the printing process, two types of errors can occur and should be corrected as follows:

- Since cartridges that are filled with deionized water or ethanol are refilled and reused, they can be reported as empty by the printer even though sufficient amount of deionized water or ethanol is still left in the cartridge. Furthermore, cartridges occasionally are not recognized by the printer. If such errors occur, the cartridge is reinserted, the protocol adjusted appropriately (e.g., by performing ‘head cleaning’), and the experiment continued.
- Parts of the printer can have reached their end of life, for example the ink pad storing the ink used, e.g., in head cleaning, can be full. In principle, the ink pad can be replaced manually and the counter can be reset for the Epson Artisan 50. Since this did not work reliably, however, we used a new printer.

